# Systematic analysis of CDR contacts and sequence constraints between T cell receptor *αβ* chains

**DOI:** 10.1101/2024.05.24.595718

**Authors:** Martina Milighetti, Yuta Nagano, James Henderson, Uri Hershberg, Andreas Tiffeau-Mayer, Anne-Florence Bitbol, Benny Chain

## Abstract

The six complementarity determining regions (CDRs) of the T cell receptor (TCR) form multiple contacts with cognate peptide and major histocompatibility complex, thus determining antigen specificity. However, the contacts between the CDRs themselves are less understood. We perform a systematic study of all available TCR structures, and identify consistent patterns of intra- and inter-chain CDR contacts. We further show that the sequences of paired TCR*α* and TCR*β* are not independent within sets of antigen-specific TCRs, for most epitopes. We quantify this sequence restriction using a mutual information framework. Co-evolution models can achieve some *de novo* prediction of TCR*α*/TCR*β* pairing, without using a training set of known pairs. The conserved pattern of CDR amino acid contacts, and the mutual sequence constraints between antigen-specific sets of T cell receptor *α* and *β* chains could play an important role in shaping the antigen-specific T cell repertoire.

## Introduction

The *αβ* T cell receptor (TCR) is a heterodimer which recognises peptides bound to major histocompatibility complex (MHC) molecules (pMHC). Each TCR chain is generated by V(D)J recombination, which generates an enormous diversity of TCR sequences (Davis & Bjorkman, 1988; Keşmir et al., 2000; Mora & Walczak, 2019). The complementarity-determining regions (CDRs or CDR1, CDR2 and CDR3) are the most variable regions of each TCR chain, and make contact with the cognate pMHC. CDR1 and CDR2 are encoded by the germline V gene, while CDR3 spans the recombination junction between V, J and D genes and is consequently the most diverse (Schwartz & Hershberg, 2013). TCRs have a conserved docking mode on pMHC, with CDR3 making the most contacts with the peptide, whilst CDR1 and CDR2 mostly contact the MHC (Szeto et al., 2020; Milighetti et al., 2021). Notably, thousands of TCRs of different sequence can bind to the same pMHC (Dash et al., 2017; Glanville et al., 2017) and a single TCR may bind to many pMHCs (Sewell, 2012).

The extent to which TCR diversity is constrained by TCR*αβ* interactions remains unclear. At a whole repertoire level, only very weak or no constraints on pairing of TCR*α* and TCR*β* sequences have been reported (Dupic et al., 2019; Yu et al., 2019; Shcherbinin et al., 2020). Shcherbinin et al., 2020 detected no constraints on TCR*αβ* pairing in PBMCs or in epitope-specific repertoires, but showed that the residues at the *α/β* interface can affect the relative orientation of the two TCR chains. However, paired TCR*αβ* have been shown to carry more information about epitope specificity than each chain independently (Carter et al., 2019; Springer et al., 2021; Mayer & Callan, 2023; Henderson et al., 2024). Experimental studies have suggested that interactions between CDR loops might be important for epitope specificity. For instance, McBeth et al., 2008 showed that residues in the CDR3s comprise over 30% of the *α/β* interface, and affect the inter-domain angle between the two TCR chains and therefore pMHC binding. Moreover, single mutations in CDR1 and CDR2 can lead to loss of antigen binding (Gras et al., 2010), but the effect of mutation is dependent on the CDR3 context (Stadinski et al., 2014). Therefore, whilst there are few or no constraints on which TCR*α* can pair with which TCR*β* at a protein-folding level, thymic and post-thymic antigen selection may restrict *α*/*β* pairing and interactions between the CDR loops may influence antigen binding. These interactions have the potential to impact TCR affinity or cross-reactivity, and may therefore constitute an important consideration in the design and optimisation of TCR-engineered T cells (Campillo-Davo et al., 2020; Baulu et al., 2023) or soluble TCRs (Robinson et al., 2021) for therapeutic applications. Indeed, amino acids at the interface between TCR*α* and TCR*β* can modulate expression and stability of transduced TCRs (Thomas et al., 2019), but their impact on TCR affinity and specificity remains to be resolved.

In this study, we systematically document the intra-chain and inter-chain interactions that the hypervariable loops make with each other. We then explore global sequence constraints on TCR*α* and TCR*β* pairing, in the context of antigen-specificity. We quantify the constraint using a mutual information (MI) framework, and use co-evolution mutual information-based models to predict pairing.

## Methods

### Analysis of existing crystal structures

A list of existing crystal structures for TCRs and TCR-pMHC complexes was obtained from The Structural T-Cell Receptor Database (STCRDab, Leem et al., 2018, downloaded on 27^th^ February 2023). The dataset was curated to include only *αβ*TCRs, both mouse and human. Structures annotated as bound for which the IMGT-renumbered file available from STCRDab had missing epitope information were removed, as well as epitopes with special groups and TCRs binding superantigens and enterotoxins. Structures with reverse orientation were analysed separately. The complete set included 509 structures. Where multiple structures were available from a single PDB, the structure with the most complete chain combination was chosen, reducing the number to 340 structures. The IMGT-renumbered sequence was then extracted from each PDB file, and TCRs with identical sequences for both TCR*α* and TCR*β* chains were identified. Using the sequence information, we retained a set of 157 bound and 78 unbound receptors, each set containing only receptors with a non-identical sequence.

Structures were visualised using Pymol Open Source v2.5.0 (Schrödinger, LLC, 2015). To calculate intra-chain and inter-chain distances, we calculated the distance between each pair of non-H atoms in the two residues under consideration using the BioPDB package in Python v3.11 (Hamelryck & Manderick, 2003). We then took the minimum distance between any two atoms.

Contacts were defined as a minimum distance *<* 5Å between two residues. A number of alternative thresholds were evaluated and 5Å was chosen, as the average number of contacts between CDRs forms an elbow around the 5Å threshold (Figure S1).

For the epitopes analysed, we retrieved the crystal structures of pMHC complexes from the Protein DataBank (PDB). Solvent-accessible surface area (SASA) for each epitope was calculated using Pymol Open Source v2.5.0 (Schrödinger, LLC, 2015) using command get_area on the peptide chain with dot_solvent 1 and solvent radius 1.4 Å on a representative structure for each epitope (Table 1).

**Table 1:**
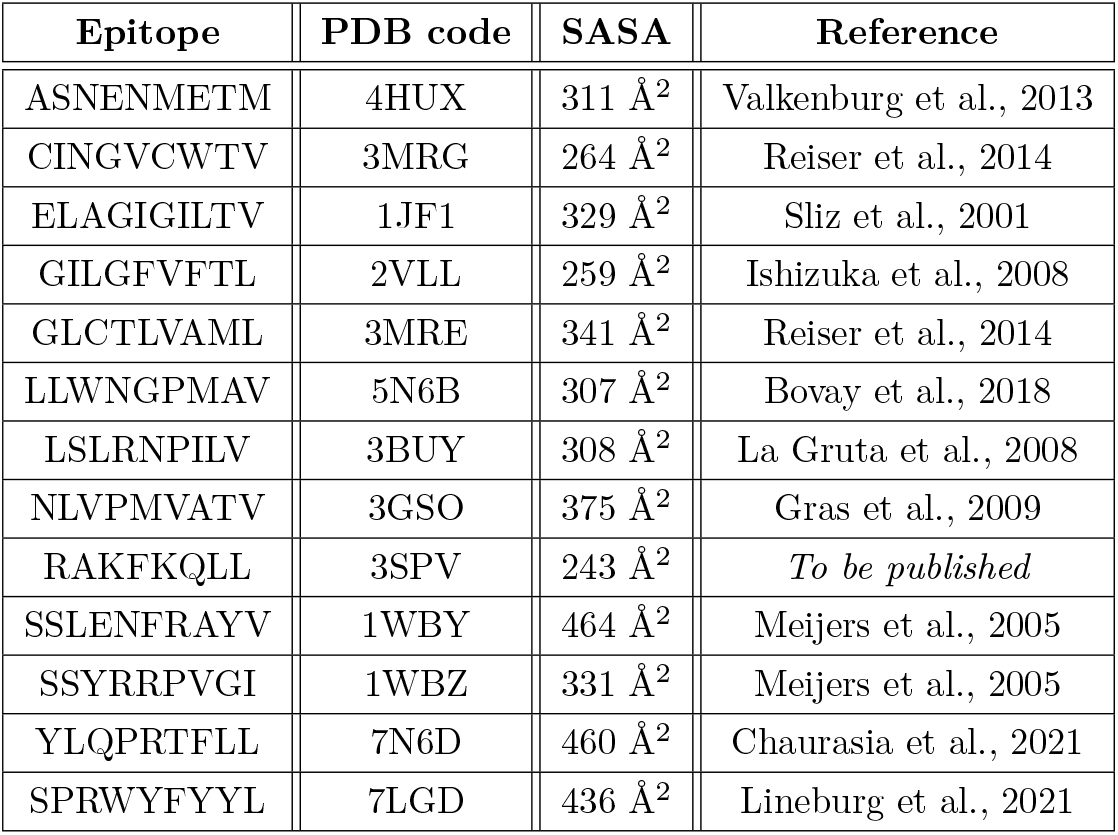
List of epitopes for which a PDB structure is available and calculated solvent-accessible surface area.

### Paired-chain datasets

The entire VDJDb database (Goncharov et al., 2022) was exported on 3^rd^ February 2023. It was refined to only include TCRs for which both TCR*α* and TCR*β* chains are available, and to include unique TCR-epitope pairs. All epitopes for which fewer than 100 or more than 10,000 sequences are available were removed from the set. stitchr (Heather et al., 2022) was used to obtain full chain sequences for each V/CDR3/J annotation for both mouse and human TCRs, and ANARCI (Dunbar & Deane, 2016) was used to obtain CDR3 sequences from each chain, renumbered to include standard IMGT gaps (Lefranc et al., 2003). The final set of sequences was used for all analysis as described in the main text (Table S1).

Pre-processed data from Tanno et al., 2020 was obtained from the authors, and further processed as in Mayer and Callan, 2023. The clonotypes from sample A1, sorted naïve cells were used as a control unselected repertoire.

We also downloaded all human data from The Observed T Cell Receptor Space (OTS) database (Raybould et al., 2024). We removed datasets which reported on *γδ* T cells, thymocytes or MAIT cells, and runs that had less than *<* 100 sequences.

To ensure that CDR3 sequences were all the same length for MI calculations, sequences longer than 19 residues were discarded (Figure S2), corresponding to *<* 1% (67 of 9852) of all CDR3*αβ*. Sequences shorter than 19 were padded by addition of gaps at position 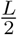 (where *L* is the length of the CDR3 sequence).

### Sequence clustering based on triplet similarity

Sequence similarity between CDR3s was assessed by normalised triplet similarity as previously described (Joshi et al., 2019). Briefly, each CDR3 is decomposed into sets of overlapping triplets. The number of triplets shared between two CDR3s is counted and normalised by 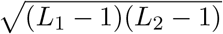, where *L*_1_ and *L*_2_ are the lengths of the two CDR3s. Triplet similarity is calculated by using the stringdot function (norm = TRUE) of the kernlab package (Karatzoglou et al., 2004). The similarity function is called in Python by using package rpy2 (https://rpy2.github.io/) and networks are plotted using python-igraph (https://python.igraph.org/en/stable/, Csárdi & Nepusz, 2006).

Triplet similarity was calculated on CDR3 sequences which did not include gaps. Sequence similarity graphs were defined by thresholding triplet similarity at a value of 0.76 and 0.72 for *α* and *β* respectively. The thresholds were set such that 99.99% of pairwise distances between CDR3 sequences not recognising the same epitope are below the threshold.

### Calculating mutual information between TCR regions

To quantify how much restriction one region of the TCR poses on another, we calculated the MI between different regions. MI is a statistical measure of the dependence of two random variables, i.e. of how much information each variable carries about the other. The MI between two regions where variability is moderate, such as V or J genes, can readily be estimated by using the observed frequencies of each gene in the set, together with their observed frequencies of co-occurrence. However, estimating the MI between one such region and a CDR3 sequence, or between two CDR3 sequences, is not as straightforward. Indeed, most sequences occur only once (Table S1) and the data is thus insufficient to estimate the entropy of CDR3 sequences as a whole (and thus the MI involving them). To circumvent this issue, we focused on pairwise approximations of the CDR3 MI, where the amino acid sequence of the CDR3 region is considered one residue at a time. The MI between a region of moderate variability and each single residue position of the CDR3 region was calculated, and then summed to obtain a pairwise approximation of the total MI between these two regions. This is an approach that has been successfully used to study protein-protein interactions and co-evolution (Dunn et al., 2008; Skerker et al., 2008; Bitbol, 2018; for an alternative approach using unbiased estimators for Renyi-Simpson entropies see Tiffeau-Mayer, 2023 and Henderson et al., 2024). Specifically, the MI between a moderately variable region *X* and a CDR3 sequence *S* was approximated by:

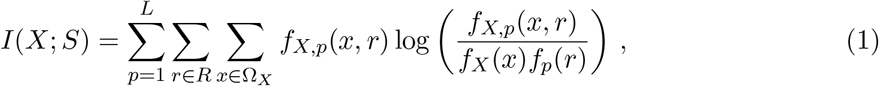

where *f* denotes a frequency observed in the data, *L* is the length of the CDR3 amino acid sequence of interest (we enforced *L* = 19 as described above), *r* is a residue at position *p* in sequence *S, R* is the set of possible residues (i.e. the 20 amino acids plus a character for a gap in alignment), and *X* is the random variable describing the moderately variable region of interest (e.g. the V gene), taking values *x* in an ensemble Ω_*X*_. Similarly, a pairwise approximation of the MI between two CDR3 regions *T* and *S* was calculated by summing over all pairs of residue positions involving one position in each of these two regions:

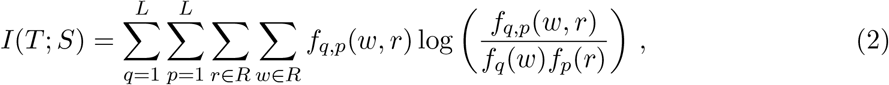

where *w* is a residue at position *q* in sequence *T* (while *r* is a residue at position *p* in sequence *S*, as before).

Since plug-in estimates of MI are biased for small sample sizes (Nemenman et al., 2002; Archer et al., 2014), the ‘real’ value of the MI was extrapolated by taking sequential subsamples of size *N* [25, 35, 50, 80, 100, 150, 200, 300, 500, 1000, 1500, 2000, 2500, 3000, 5000, 10000, 15000] until sample size was reached. A regression line was then fitted to the MI versus 1/*N* at each subsample and the y-intercept was used as the MI estimate (lim_*N→∞*_) (Strong et al., 1998; Panzeri et al., 2007). The procedure was repeated for both the real set and a shuffled version of the same set, where one variable is kept constant, whilst the second variable is randomly shuffled, so that the background entropy remains the same but the relationship between the two variables is broken. To account for batch effects that might be dependent on sequencing technology or study design, we perform the shuffle only within the sequences belonging to a single study (thus maintaining any batch effects on the signal). In order to allow within study randomisation we restricted the analysis to studies which contain at least 3 sequences for all MI analyses. Each subsampling was repeated 10 times. Figure S3 shows the estimation for epitope GLCTLVAML as an example.

MI was calculated using scikit-learn v1.1.3 (Pedregosa et al., 2012). To reduce the diversity in V genes and J genes, tidytcells v1.6.0 (Nagano & Chain, 2023) was used to standardise to the gene level, removing allele information. We further dropped duplicates in TRAV-CDR3A-TRAJ / TRBV-CDR3B-TRBJ pairs within each epitope repertoire after gene standardisation to avoid inflating the estimated MI.

### Calculation of effective set size

We define effective set size as the number of distinct sequences present in a repertoire, accounting for repeated or similar sequences. It is calculated as in Weigt et al., 2009 and Bitbol et al., 2016. Briefly, for each sequence (*S*) in a set of size *N* the number of sequence neighbours within a similarity threshold are counted (*m*_*S*_). *S* is then given a weight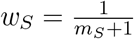. The total effective set size can finally be calculated as:

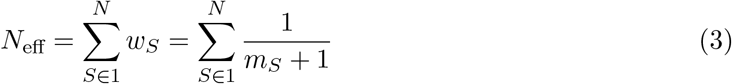

If all sequences are distinct and have no neighbours, *N*_eff_ = *N*, whilst if all sequences are neighbours of all other sequences *N*_eff_ = 1. The diversity is calculated by Hamming distance on the padded CDR3 sequences, excluding the first 3 and the last 3 amino acids of the CDR3. We set the threshold of similarity to be up to Hamming distance 1 for CDR3*α* and CDR3*β*, and up to Hamming distance 2 for CDR3*αβ* (calculated as a sum of the distance over the two chains). The effective set size *N*_eff_ is then normalised by total repertoire size *N* for comparison across repertoires. The effective set size was not correlated to total repertoire size (Figure S4).

### Mutual information iterative pairing algorithm (MI-IPA)

The mutual information-based iterative pairing algorithm (MI-IPA) was described in Bitbol, 2018. The algorithm aims to correctly match pairs of interacting proteins from two input lists based on the MI between the respective amino acid sequences, by permuting the pairing between the two lists to approximately maximise MI. The method exploits the observation that, in families of evolutionarily related interacting proteins, the interacting contact regions between the proteins will co-evolve, such that a change in sequence of one member of the pair will constrain the possible compensatory changes in the other member of the pair, in order to preserve the required functional interaction. We adapted the method to pair a set of unpaired TCR*α*/*β* sequences and implemented it in Python v3.11.0. Correct pairings were known but blinded to the algorithm.

The MI-IPA is run on each epitope-specific repertoire from VDJDb, without using a training set of known pairs (as would be the case if no information about pairing was available) and inputting the CDR3 sequences only. For epitopes that have *>* 1000 sequences, 5 subsamples of 700 sequences were generated and paired, to reduce computational time. The original implementation of the MI-IPA used species to reduce the combinatorial number of assignments to be evaluated. As species are not available in this case, we used study and individual information as available from VDJDb as a substitute. As repeats are present both in the TCR*α* and TCR*β*, randomness was added into the model at the assignment stage. To account for this, as well as the influence of the initial random pairing, we ran the model 10 times in each condition.

Parameters *θ* (sequence diversity threshold) and *λ* (pseudocount) were defined as in Bitbol, 2018 and used to correct the frequency calculation for MI estimation. Step size and confidence calculation were also optimised for the dataset. We compared the Hungarian scoring method, a greedy assignment based on confidence calculation, and a ‘no confidence’ scenario, where the greedy assignment is performed directly on the scores assigned to each pair, without calculating a confidence score. These parameters are optimised on epitopes GLCTLVAML and YLQPRTFLL (Figure S5). We run the MI-IPA with diversity threshold *θ* = 0.6, pseudocount *λ* = 0.6, step size of 6 sequences and no confidence calculation. Equivalent settings but with *λ* = 1 were used as a negative control yielding the chance expectation (as *λ* = 1 prevents the algorithm from learning).

### Graph alignment (GA) algorithm

The graph alignment algorithm (GA) aims to align two sequence similarity networks built on the lists of sequences to be paired (Bradde et al., 2010). The method assumes that two co-evolving protein families will have similar phylogenetic trees (in our use-case, similar similarity graphs), and thus tries to align the similarity of the two trees to pair the family members. First, all pairwise *α − α* and *β − β* CDR3 distances in each input list are calculated. The distances are then used to calculate the two similarity networks and the generated graphs are aligned by trying to maximise the number of overlapping edges. Gandarilla-Pérez et al., 2023 recently published an implementation of the GA, with code freely available at https://github.com/carlosgandarilla/GA-IPA. The published code was retrieved and integrated in our pipeline. No amendments were made to the code that performs the alignment. To allow integration between the GA, which was coded in Julia v1.8.5, and the existing Python v3.11.0 pipeline we used PyJulia v0.6.0 (https://pyjulia.readthedocs.io/en/latest/).

Four different distance measures were used: (1) Levenshtein (or edit) distance, (2) Weighted Levenshtein distance (where substitutions are weighted 1 and gaps are weighted 1 + ln(4)), (3) Triplet distance (calculated as 1-triplet similarity, as described above), (4) TCRdist (Dash et al., 2017) calculated on the CDR3 sequence only. Levenshtein and weighted Levenshtein distance are implemented in rapidfuzz (https://pypi.org/project/rapidfuzz/ Bachmann, 2024), whilst TCRdist is calculated using the implementation available in package pwseqdist (https://github.com/agartland/pwseqdist), using the parameters specified within tcrdist3 (https://tcrdist3.readthedocs.io/en/latest/, Mayer-Blackwell et al., 2021).

As for the MI-IPA, the GA was run on each epitope separately. Epitopes with *>* 1000 sequences were subsampled 5 times to 700 sequences to reduce computational time. The same two epitopes as for MI-IPA, GLCTLVAML and YLQPRTFLL, were used to optimise the distance metric and the number of neighbours (*k*, defined as in the original method) to achieve the best pairing (Figure S6). The GA was run using Levenshtein distance and *k* = 20.

### Combining GA and MI-IPA (GA+MI-IPA)

We combined the GA and MI-IPA as proposed by Gandarilla-Pérez et al., 2023. In the original implementation, the GA was used to calculate a first pairing for the sequences of interest. The stable assignments (i.e. the pairs that were consistently assigned over many iterations of the GA) were then used as a golden set for the MI-IPA (i.e. a training set of pairs that are used to initialise the MI-IPA and kept as they are throughout the IPA). Here, we explored three different ways of combining GA with MI-IPA:

(1) the most stable pairs (selected *≥* 95% of the time, or the top 5 pairs, whichever is largest) from the GA are used as a golden set for the MI-IPA and removed from the set of pairs to be paired by the MI-IPA (as in Gandarilla-Pérez et al., 2023);
(2) the most stable pairs (selected *≥* 95% of the time, or the top 5 pairs, whichever is largest) from the GA are used as a golden set for the MI-IPA but they are not removed from the set of pairs that the MI-IPA tries to assign. This allows the MI-IPA to correct the pairs in the final assignments;
(3) the complete consensus assignment from the GA (i.e. the TCR*β* that is most often assigned for each TCR*α* across 100 repeats of the GA) is used as an input training set to the MI-IPA, but all *α* and all *β* are also available for re-pairing by the MI-IPA.

Note that when multiple *β* are chosen the same number of times for an *α* in the GA, the *β* included in the stable assignment is chosen at random. The final assignments for all pairs are then extracted and evaluated against the known pairs.

As for the MI-IPA and GA, also the GA+MI-IPA was run on each epitope separately, using 5 subsamples of 700 sequences for epitopes with *>* 1000 sequences. Epitopes GLCTLVAML and YLQPRTFLL were used to find the optimal combination of GA+MI-IPA (Figure S7). We ran the GA+MI-IPA with the combination regime described in (2) above.

### Calculation of precision

To quantitatively compare pairing algorithms, we calculated the proportion of predicted pairs that are correct (precision). Note that there are repeats in the *α* and *β* chain sets which will influence this calculation (see example in Figure S8A, B). Importantly, the calculation of precision takes into account information about which individual repertoire a TCR sequence is found in. Therefore, a pair must be correct and must be found in the correct individual to be called correctly assigned. For instance, if a pair belonging to individual A but not individual B is predicted for individual B, it is marked as incorrect (Figure S8A, C).

### Benchmarking the pairing models

To benchmark the performance of MI-IPA, GA and GA+MI-IPA, we predicted the pairing using TULIP (Meynard-Piganeau et al., 2024) and SCEPTR (Nagano et al., 2025). TULIP prediction was performed using the available pre-trained model. For each epitope, we generated all possible CDR3*α*-CDR3*β* pairs available from each individual. We then predicted the probability of binding of each pair against the epitope. We then assigned *αβ* pairs with an algorithm similar to that used for the MI-IPA. Briefly, for each individual, we selected the most likely pair (lowest score). We then removed that *α* and the *β* sequence from the set, assigned the best remaining pair and so on. We calculated precision as above.

To assign pairing with SCEPTR, we embedded the *α* and the *β* chains separately using the pre-trained default model. Since the training is performed by masking sections of the TCR, we expect that *αβ* paired chains may have similar embeddings. Therefore, we calculated the Euclidean distance between all *α* and all *β* chains from one individual for each epitope and then assigned pairs as above (starting with the pair with the smallest distance) and then calculated precision as previously defined. SCEPTR was run both with its default model (trained on paired data) and with a model trained on synthetic data, in which the pairing in the training set was random (SCEPTR - synthetic).

## Results

### Interactions between CDR loops in existing TCR structures

To document CDR interactions, we examined the TCR-pMHC structures available from the The Structural T-Cell Receptor Database (Leem et al., 2018, STCRDab). Observation of 4 representative TCRs (Figure 1) shows that CDR loops form a continuous contact surface that binds pMHC. There are both intra- (between two CDR residues on the same chain) and interchain (between one residue on the *α* and one residue on the *β* chain) contacts (middle and right panels for each TCR) observed both in the bound (right panel, indicated with a *) and in the unbound (middle panel) TCR structures, although the number of contacts varies between the ligated and unligated version.

**Figure 1.**
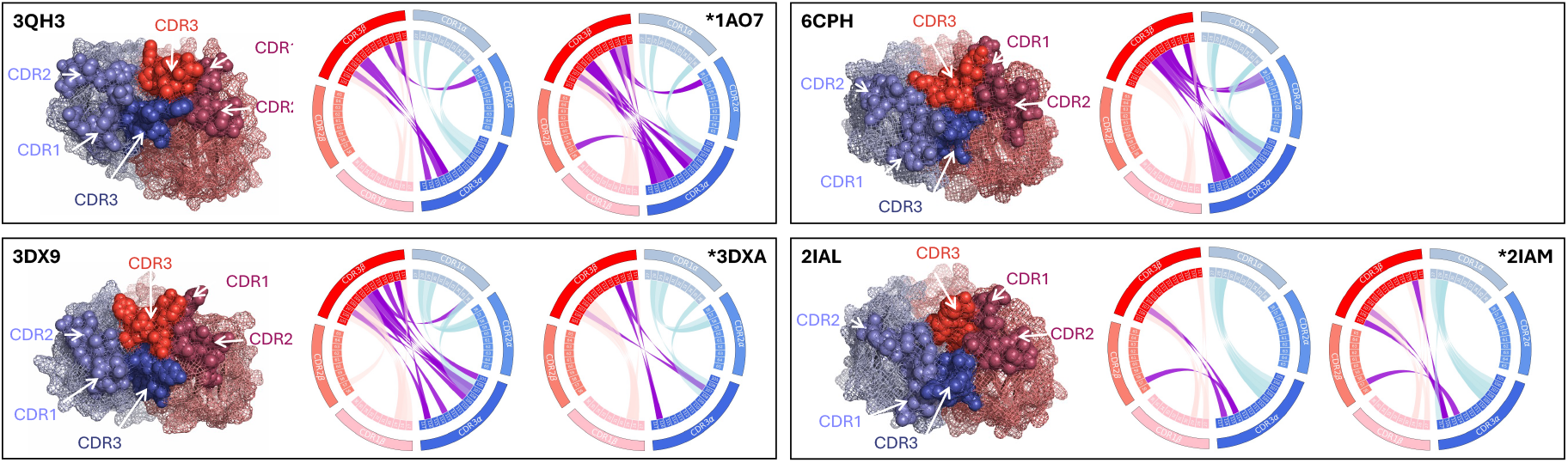
CDR loops come together to form the TCR binding site in four example TCRs. Positioning of CDR loops of 4 representative TCRs (2 recognise epitope on class I MHC, 3QH3 and 3DX9; 2 recognise epitope on class II MHC, 6CPH and 2IAL). The TCR*α* chain is in shades of blue, whilst the TCR*β* is in shades of red. CDR1, 2 and 3 for each chain are represented with spheres and annotated on the figure. Mesh shows the rest of the TCR structure. On the right of each structure, the circle plot shows contacts between the CDR loops (defined as minimum distance between any two non-H atoms *<* 5Å) for the same TCR in both the unbound (left) and bound (right, indicated with a *\**) structure. Light blue: contacts between loops on TCR*α*; light red: contacts between loops on TCR*β*; purple: contacts between the two chains. 3QH3: Scott et al., 2011, 1AO7: Garboczi et al., 1996, 3DX9 and 3DXA: Archbold et al., 2009, 6CPH: Galperin et al., 2018, 2IAL and 2IAM: Deng et al., 2007.

To more systematically map the interactions between TCR residues, we first measured the intra-chain distances between each residue and all other residues on the same chain for the four example unligated structures (Figure 2A and B for TCR*α* and *β*, respectively). Consistent with Figure 1, we see extensive intra-chain contacts, especially between CDR1 and CDR3 and CDR1 and CDR2, both in the TCR*α* and in the TCR*β* chain. Remarkably, the patterns are consistent across the four structures and in both chains. We extended this analysis to 157 unique pMHC bound TCRs (Figure 2C, D and Figure S9B, C for comparison between class I and class II bound TCRs) and 78 unique unbound receptors (Figure S9A). Contact is defined as minimum distance between two non-H atoms on two residues *<* 5 Å. The patterns observed in the 4 example structures are generalisable and observed in TCRs both in the unligated and ligated form (class I and class II): CDR1 forms extensive contacts with CDR3, and CDR2 in turn forms contacts with CDR1. We also observe conserved contacts between the first half of CDR2 and the first half of CDR3. Strikingly, all 340 PDB files analysed show contacts between CDR1 and CDR3 on both chains (Figure S11, left). We also observe similar patterns in TCRs that bind MHC with reversed polarity (Figure S9D).

**Figure 2.**
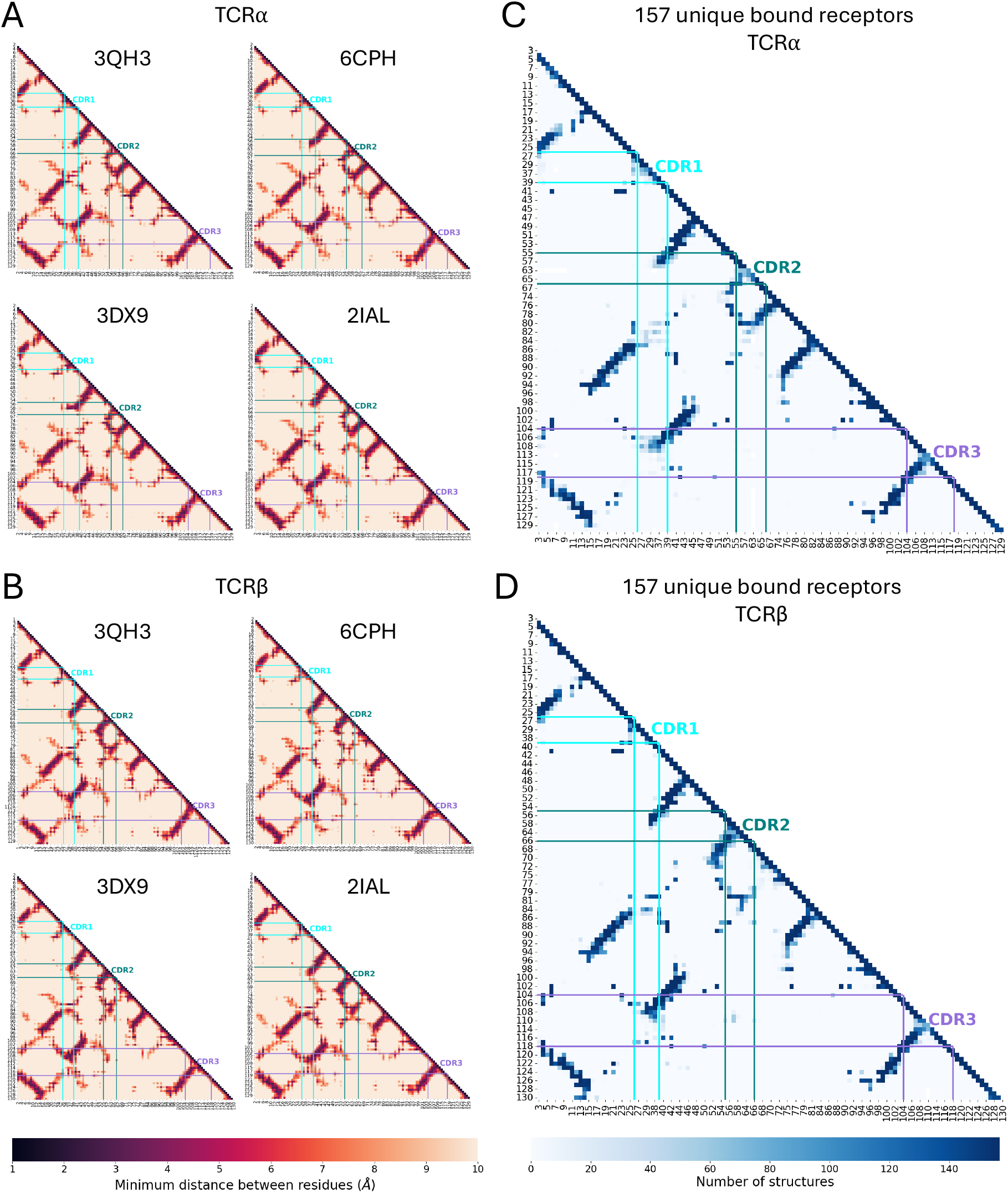
Conserved intra-chain interactions are observed in TCR structures. **A, B)** Intra-chain contact maps for the unligated structures in Figure 1 for the TCR *α* and *β* chain, respectively. The distance (Å, colour bar) is calculated between all pairs of atoms in the two residues, and the minimum distance is shown. **C, D)** Intra-chain contact maps were calculated for the TCR *α* and *β* chain (respectively) of 157 unique bound TCRs. A contact was defined as distance *<* 5Å. The heatmap shows the number of structures that are found to make contact at each pair of positions. Only positions present in *>* 75% of structures according to the IMGT numbering scheme are shown. Only the bottom half of each distance matrix is shown as it is symmetrical across the diagonal. In each heatmap, the start and end positions of CDR1, 2 and 3 are highlighted.

We then asked whether interactions can be observed between a CDR in one chain and a CDR in the other chain (inter-chain contacts). Figure 3A shows the inter-chain distances for the four unligated TCRs in Figure 1. Again, the distance patterns are conserved across the four TCRs, both in the conserved framework regions and in the CDR loops. Specifically, framework region 2 (FR2), found between CDR1 and CDR2, makes conserved contacts with FR2 on the other chain, as well as contacts with the beginning and end of the CDR3, where the CDR3 sequence is least variable (the diamond-shaped pattern). We speculate these are conserved structural contacts that allow proper formation of the TCR heterodimer. As well as these, inter-chain contacts are observed between CDR1 and 2 and the CDR3 on the paired chain. Remarkably, we observe a high number of contacts between the two CDR3s, whilst no contacts are observed between the germline loops across the two chains. Notably, the two CDR3s form a X-shaped pattern: the interactions here involve the variable junctional regions, but no interactions are observed between the variable regions and the conserved segments on the loop (the opposite of the diamond pattern above). We extended these observations to the whole set of bound and unbound TCRs (Figure 3B and Figure S10A-C). The diamond-shaped pattern between FR2 and CDR3*α*/*β* contacts is conserved across most structures, as are the interactions between the CDR3s. Notably, these patterns can also be observed in TCRs which bind with reverse orientation (Figure S10D). As for intra-chain contacts, all 340 PDB files analysed show inter-chain contacts between CDR3s (Figure S11, right). Overall, these results suggest that the CDR loops come into close proximity with one another to form the antigen binding site, even prior to pMHC binding. We predict that interactions between the CDRs could stabilise the conformation of the loops and hence impact the kinetics of binding, but further structural and functional studies will be required to test the prediction.

**Figure 3.**
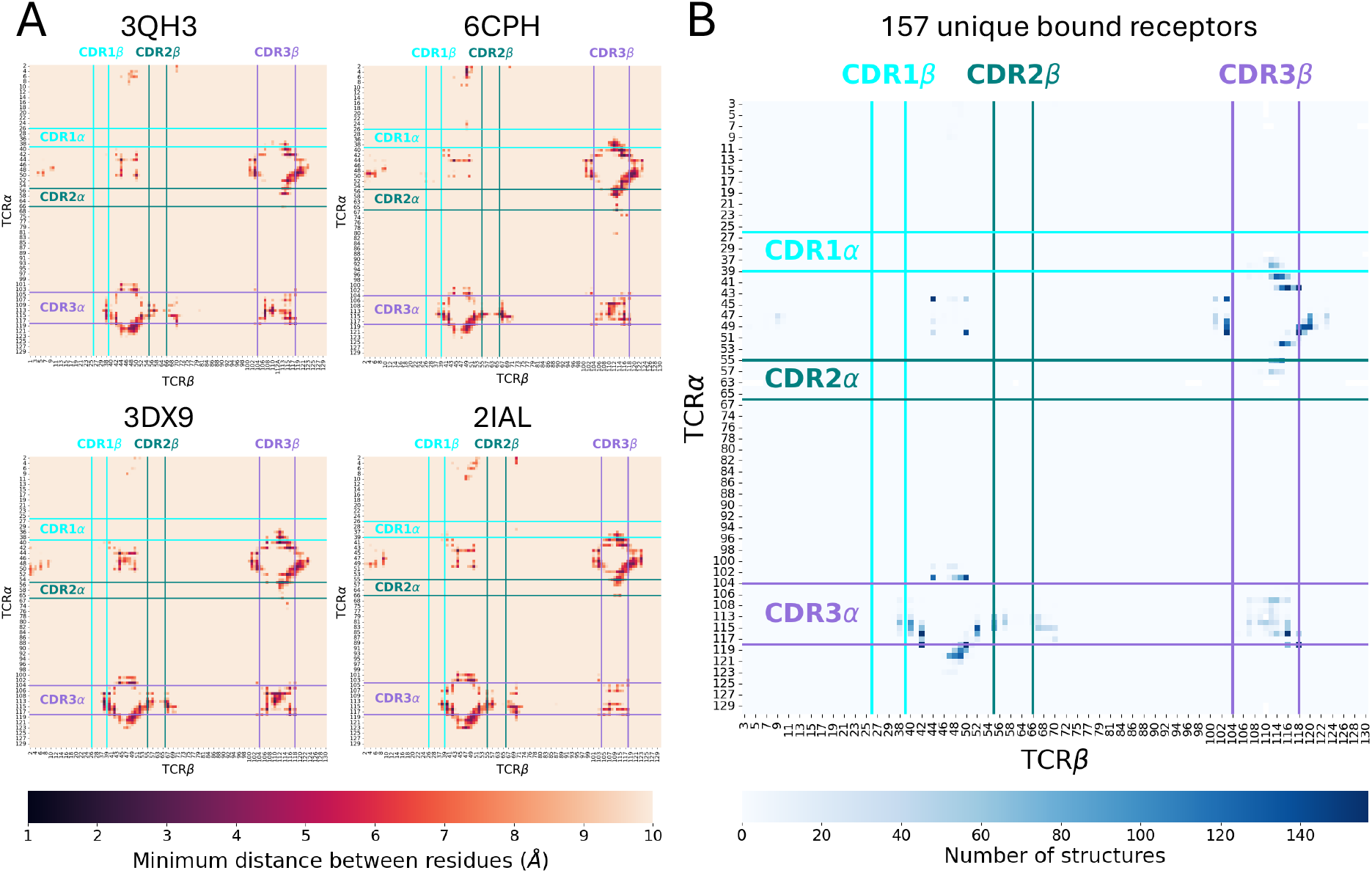
Conserved inter-chain interactions are observed in TCR structures. **A)** Inter-chain (TCR*α* to TCR*β*) contact maps for the four structures in Figure 1. The distance (Å, colour bar) is calculated between all pairs of atoms in the two residues, and the minimum distance is shown. **B)** Inter-chain contact maps were calculated for 157 unique bound TCRs. A contact was defined as distance *<* 5Å. The heatmap shows the number of structures that are found to make contact at each pair of positions. Only positions present in *>* 75% of structures according to the IMGT numbering scheme are shown. In each heatmap, the start and end positions of CDR1, 2 and 3 along the sequence are highlighted.

### A footprint of inter-chain interactions on TCR*α* and *β* paired sequences

We reasoned that the close spatial arrangement of the CDR regions in the binding region of the folded TCR may impose some constraints on the pairing between TCR*αβ* amino acid sequences recognising a given antigen. Thus, within an antigen-specific set of TCRs, the sequence of one TCR chain may restrict the allowed sequence on the other chain, due to direct contacts or enforced conformations. One testable prediction of this hypothesis is that a set of similar antigen-specific TCR*α* should bind to a set of similar TCR*β*, and vice versa. Furthermore, this constraint should be specific to antigen-specific repertoires, and very low in background repertoires.

To test this hypothesis, we retrieved paired-chain data from the VDJDb and analysed each epitope-specific repertoire (Bagaev et al., 2020; Goncharov et al., 2022). We clustered all CDR3*α* and all CDR3*β* sequences based on triplet similarity (see Figure 4A for a representative example). We observed no one-to-one relationship between clusters of *α* sequences and clusters of *β* sequences. However, within a single cluster of CDR3*α* sequences, the similarity of the paired CDR3*β* is higher than size-matched random subsamples of CDR3*β* sequences from that same repertoire (Figure 4B), and similarly for CDR3*β* clusters. Notably, all but one *α* clusters (cluster 139) and 5 of 7 *β* clusters (clusters 0, 1, 6, 89 and 94) from epitope GLCTLVAML show higher similarity in paired sequences compared to background (*β* cluster 5 shows a similar trend but does not reach statistical significance). We extended the analysis to all available epitopes within the VDJDb for which there are at least 100 paired *αβ* TCRs (Figure 4C and Figure S12). We restricted the analysis to clusters of size 3 or greater. We detected no CDR3*α* clusters of size 3 or greater in epitopes LLWNGPMAV, SSYRRPVGI and TTDPSFLGRY and no CDR3*β* clusters of size 3 or greater in epitopes RLRAEAQVK and SPRWYFYYL. Most epitopes showed increased similarity in TCR*α* or *β* associated with clusters of the other chain (Figure S12) and indeed the pairwise similarities of CDR3*α* and CDR3*β* correlate (Figure S13). However, the increase in sequence similarity was variable, ranging from epitope GLCTLVAML where there is a clear increase in median sequence similarity in both CDR3*α* and CDR3*β*, to epitope AVFDRKSDAK where no signal is detected for either chain, and the sequence similarity even decreases for CDR3*β*. The latter suggests chain independence, i.e. TCRs binding to this particular epitope are not constrained in the CDR3*αβ* pairing.

**Figure 4.**
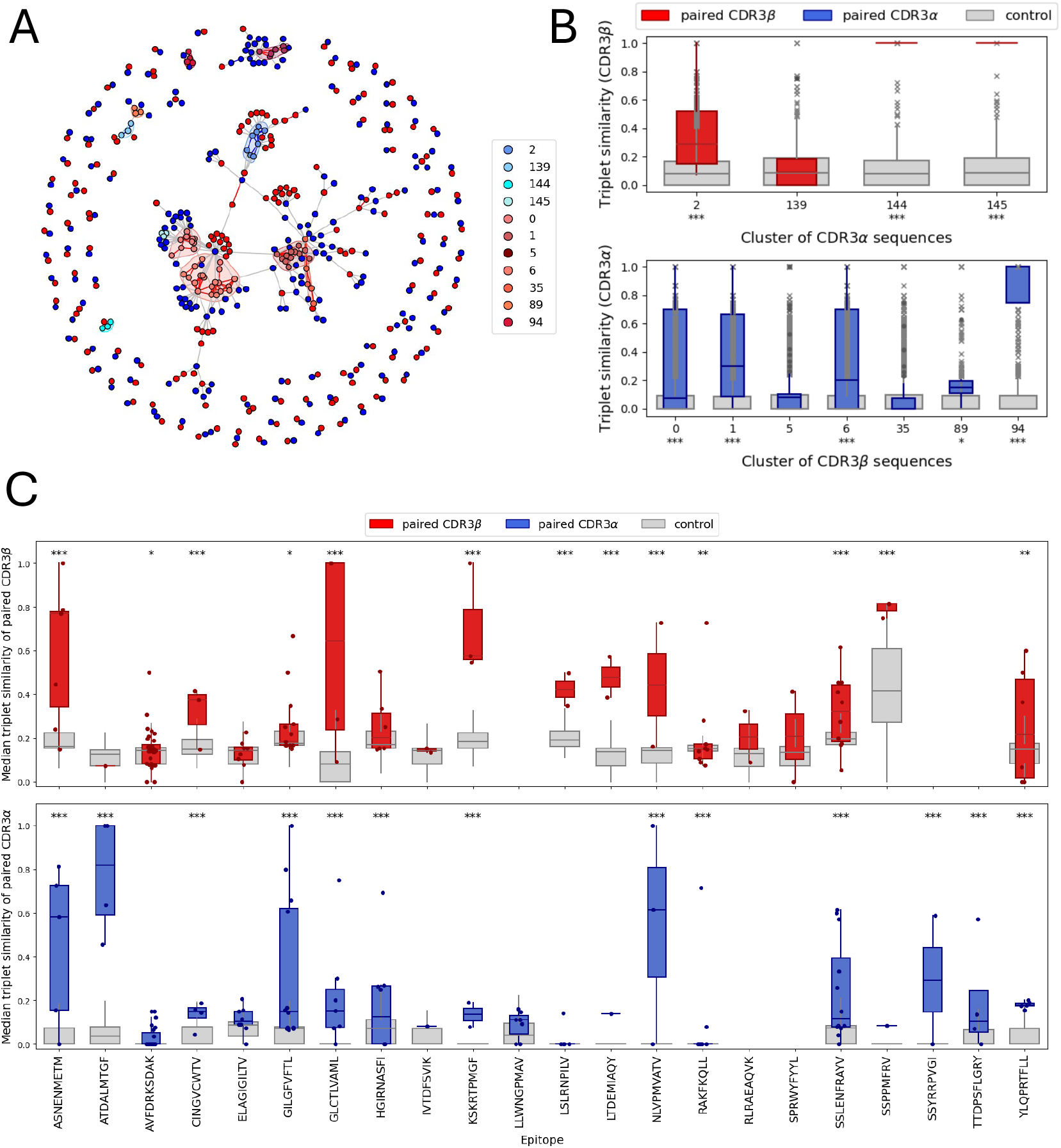
Co-clustering of CDR3*α* and *β* sequences. **A)** Each circle is a unique CDR3 sequence (*α* in blue, *β* in red). Similar *α* sequences are linked by blue edges, similar *β* sequences by red edges. Each *α* is linked to its paired *β* by a grey edge. The clusters analysed in **B** are coloured in shades of blue (*α* clusters) or red (*β* clusters) and identified by a shaded area around them. Each CDR3 sequence is represented only once, therefore the same CDR3*β* may be linked to multiple CDR3*α* (and vice versa), if it is not unique in the epitope-specific repertoire. **B)** Similarity of paired CDR3*β* when CDR3*α* are clustered (top, or vice versa - bottom) for the clusters identified in **A**. For each cluster, all pairwise similarities between the paired sequences are calculated (coloured boxplots). 100 equal-size random samples are taken as controls from the epitope-specific repertoire (grey boxplots). Only clusters of size 3 or greater are included. **C)** The analysis in **B** was repeated for 22 epitopes. Each dot represents the median for a cluster of size 3 or greater. Median similarity was also calculated for 100 controls for each cluster (grey boxes, random sequences drawn from the same epitope-specific repertoire). Columns are empty when no clusters of size 3 or larger are found. The edges of each boxplot show the quartiles, and the whiskers extend to 1.5 the interquartile range. The median of each boxplot is shown as a line. Outliers are shown as circles for real samples and crosses for the control subsamples. P-values (one-tailed t-test, comparing real to control): * < 0.05; **<0.01; ***<0.0001.

Overall, these results suggest that, within an epitope-specific repertoire, similar TCR*α* chains tend to pair with TCR*β* chains that are more similar than random, i.e. TCR*αβ* pairing is constrained.

### Quantifying the interaction between CDR loops sequences using MI

In order to generalise the quantification of interchain constraints, we re-formulated the problem in the context of information theory. We estimated the MI between TCR*α* and *β* sequences, and between V or J genes and CDR3 sequences, using a pairwise approximation and corrected for sample size and background distributions (see Methods and Figure S3). As negative control for the antigen-specific analysis, we estimated the MIs for all TCRs in VDJDb (referred to as background), as well as for a single sample of sorted naïve T cells from Tanno et al., 2020 (referred to as Tanno::A1::naïve). The former contains information about the 22 epitopes under study pooled together (and might therefore have some residual epitope-specific signal), whilst the latter should only depend on the biology of receptor formation and thymic selection, without the influence of post-thymic antigen selection.

Figure 5A shows the estimated MIs for each epitope. Since absolute MI estimation values cannot be directly compared across MI pairs with different alphabet sizes, we show these as fold change from VDJDb background. As expected, the naïve repertoire shows no increased statistical dependence from the combined VDJdb repertoire. On the other hand, most individual epitopes show some level of statistical dependence beyond background. A few epitopes, such as AVFDRKSDAK show low to no signal. The MIs for all epitopes are summarised in Figure 5B. Most inter-chain relationships show a marked increase in epitope-specific repertoires compared to background.

**Figure 5.**
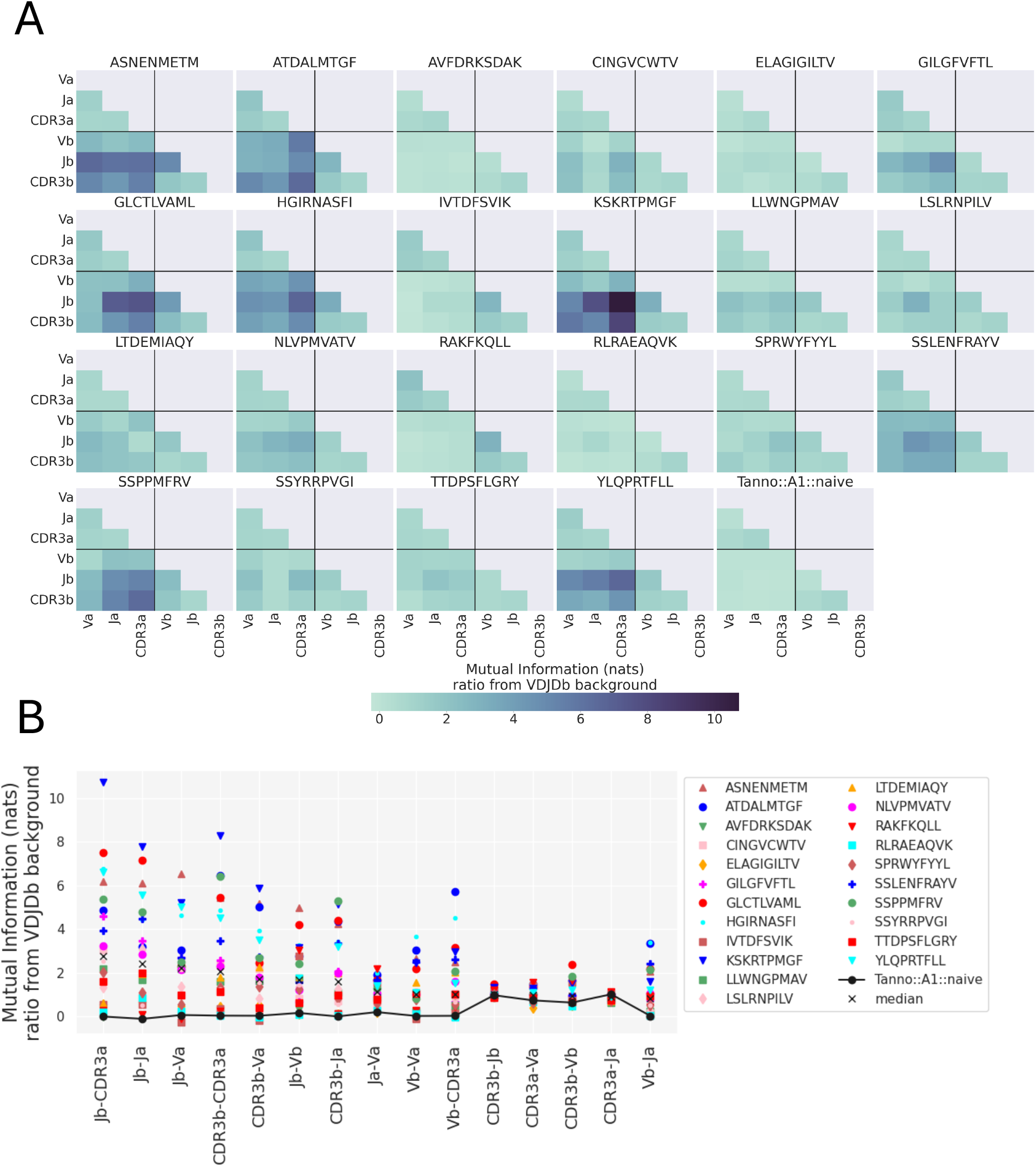
Mutual information between TCR components in epitope-specific repertoires. **A)** Mutual information (MI) was calculated between all different components of the TCR (V genes, J genes, CDR3 sequences) for each epitope-specific repertoire. In each heatmap, the top-left and bottom-right quadrants contain intra-chain MI (*α* in top-left and *β* in bottomright). The bottom-left quadrant shows the inter-chain MI. The MI was estimated for various subsamples for each epitope, and an estimate correcting for finite size effects was found for both the real set and a shuffle (Figure S3). The MI plotted here corresponds to the difference between the estimated MI for real and shuffle, divided by the MI calculated from the full VDJDb dataset. A pairwise approximation is used in cases involving CDR3 (see Methods). **B)** Summary of the plots in **A**. The x-axis is ordered according to the median MI across all epitope repertoires (indicated with an X). Each epitope is identified by a combination of colour and shape.

To understand the statistical dependence between V gene and CDR3 in more detail, we examined the estimated intra-chain MI between the V gene and each CDR3 residue (Figure 6A). As CDR3 positions 104 and 118 on both chains are constant, the MI is 0. Residues 105-107 on CDR3*α* are strongly dependent on the V gene, which reflects the overlap between V region and the start of the CDR3, and can be observed in both VDJDb and naïve background (bottom rows of the heatmap). A similar pattern can be observed with the first few residues on the CDR3*β*. However, we observed statistical dependence between the V gene and the central residues of the CDR3 for most epitopes, as well as the VDJDb background. In contrast, no statistical dependence was observed for these positions in the naïve repertoire.

**Figure 6.**
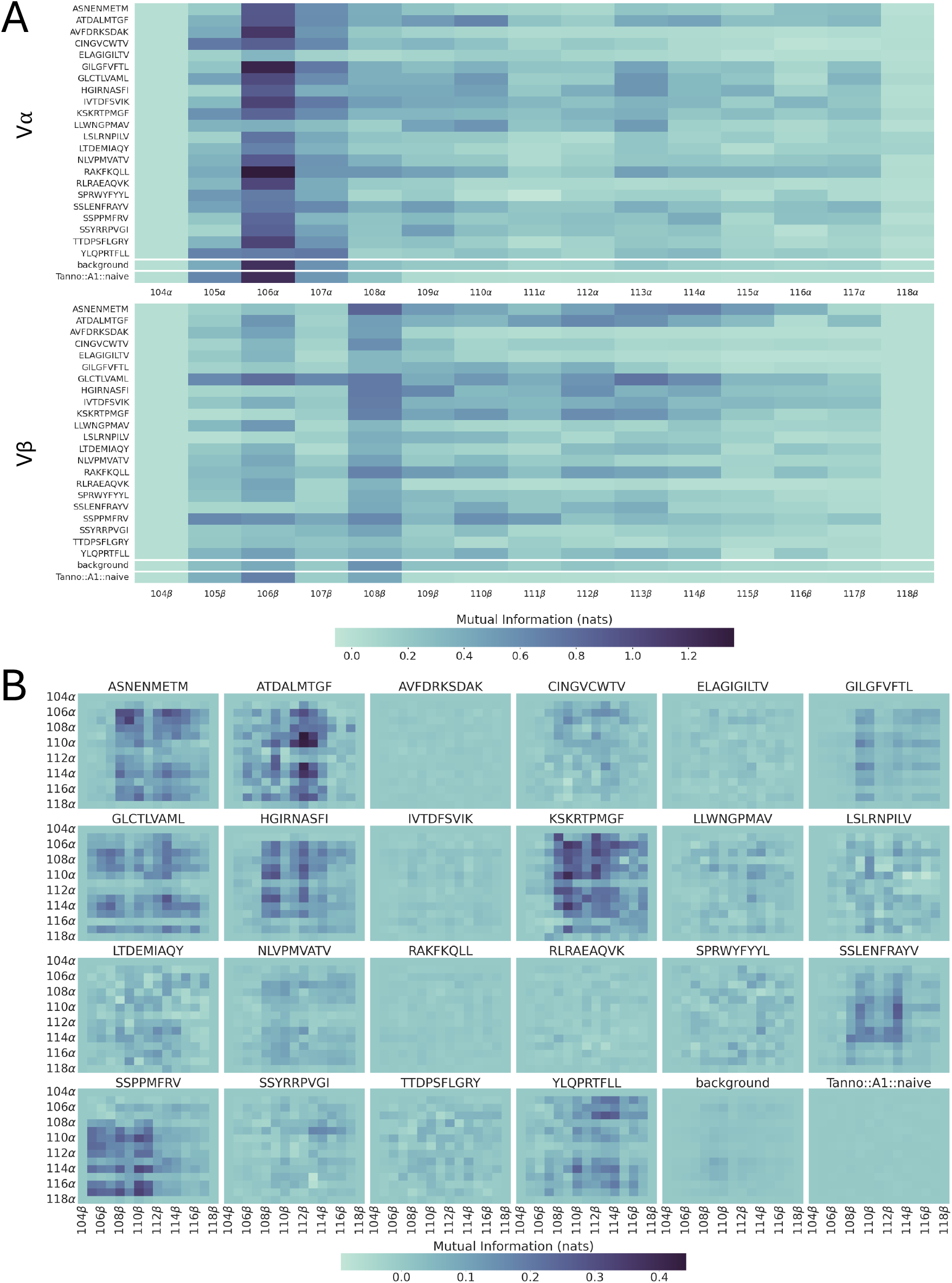
Mutual information between CDR3 and associated V gene and CDR3*α*- CDR3*β*. **A)** Mutual information (MI) between V genes and each residue position on the CDR3. Top: V*α* to CDR3*α*; bottom: V*β* to CDR3*β*. **B)** MI between each pairs of positions on CDR3*α* and CDR3*β*. The MI was calculated with pairwise approximation, extrapolation and correction for background (for both **A** and **B**, see Methods). IMGT positions between residues 111 and 112 are not shown as they are not present in most TCRs analysed.

We also examined the MI between the CDR3*α* and CDR3*β* (Figure 6B). Little or no MI is detected in the background repertoires suggesting weak restriction of CDR3 pairing at the repertoire level. In contrast, most epitopes show statistical dependence in inter-chain CDR3 interactions, with an epitope-specific signal pattern. As in all the previous analysis, epitope AVFDRKSDAK shows very low signal. Antigen-dependent TCR selection therefore commonly generates significant statistical dependence between the CDR3 sequence and V gene, as well as the CDR3*α* and *β* sequences, which we hypothesise is due to the observed structural interactions and indirect conformational effects restricting allowed chain pairing.

Since the CDR3 sequences were padded to obtain the alignment needed to calculate MI, we evaluated the effect of padding on the calculated MI. First, we correlated the proportion of CDR3 sequences that show a gap at a specific position with the average MI for that position (Figure S14) and found that positions which are more likely to have gaps tend to have lower MI. We then compared the MI estimated from sequences padded in the middle to sequences padded at the end (Figure S15). Overall, the values were well correlated between the two methods (Spearman *ρ >* 0.95) and sat close to the diagonal, with the exception of J*α*-CDR3*α* and J*β*-CDR3*β*, where the correlation was less strong, and CDR3*α*-CDR3*β*. Indeed, the J gene influences the sequence of the C terminal of the CDR3, therefore a change in alignment in this region changes the estimated MI between the two. Moreover, padding in the middle may also be slightly underestimating the MI between CDR3s by inserting gaps in the middle of the sequence, but the relative signal of all epitopes and background is maintained with both strategies. Overall, the MI calculation seems robust to the padding choice.

Finally, we wondered if the observed MI footprints observed in epitope-specific repertoires could also be observed in more polyclonal subsets, containing TCRs specific for multiple antigens. To study this, we downloaded the available datasets from the OTS database (Raybould et al., 2024) and calculated corrected MI for all the included studies (Figure S16). Consistent with the observations from VDJDb and Tanno datasets, we find that sorted memory and AIM+ cells (which presumably are enriched for antigen specific groups of TCRs) tend to have higher CDR3*α*-CDR3*β* MI, compared to unsorted and unstimulated T cells.

### Examining MI variation across epitopes

A striking feature of both the intra-chain and particularly the inter-chain statistical dependence was the variability between epitopes. Understanding the factors which drive this epitope dependence may have important implications for our ability to predict antigen specificity.

We first measured the correlation between estimated MI and epitope repertoire size, to check that our results were not caused by finite size effects. Reassuringly, we did not detect a significant correlation for most of the relationships investigated (Figures 7 and S17A).

**Figure 7.**
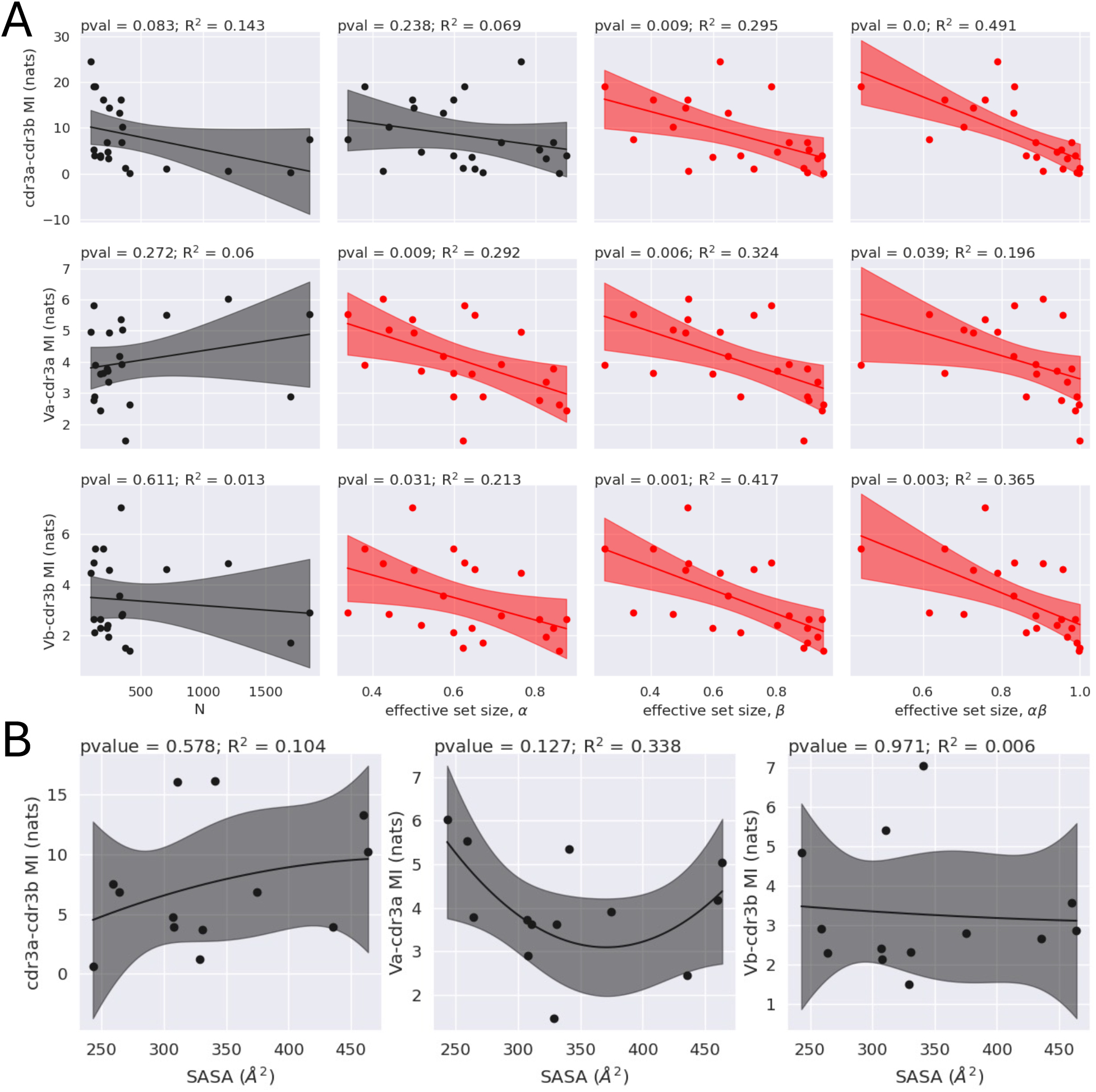
Mutual information correlates to effective set size but not peptide solvent-accessible area. **A)** Correlation of mutual information (MI) and effective set size. Effective set size was calculated by grouping all sequences with Hamming distance of *<* 2 on each chain and *<* 3 for the paired chains, normalised by total repertoire size (N). Correlation with N is also shown as a control, and no significant correlation was found in this case. All correlations with effective set size and N are shown in Figure S17. **B)** Correlation of MI and solvent-accessible surface area (SASA) for the available epitopes (Table 1). Correlations were calculated by fitting a linear regression. P-value for the F-test and *R*^2^ was calculated for each fit. Fits that have p-value<0.05 are highlighted in red. The shaded area represents the 95% CI of the fit.

Certain epitope-specific responses are characterised by usage of a specific public TCR chain which pairs with different partner chains (see for instance Zhong et al., 2007; Pogorelyy et al., 2022), or differing levels of sequence similarity (see for instance Dash et al., 2017). To evaluate the impact of similar or public TCR chains in MI estimation, we looked at the correlation between MI and the effective set size of CDR3*α*, CDR3*β* and CDR3*αβ* in each repertoire (Figure 7A and Figure S17B). We observe that repertoires dominated by a single sequence or cluster of similar sequences (a smaller set size) tend to have more MI.

We also examined whether the MI may depend on how many features the peptide residues offer for the TCR to bind, which has been proposed to correlate with repertoire diversity (Turner et al., 2005; Turner et al., 2006). We approximated the “featureness” of a peptide by the solvent-accessible surface area (SASA) of the peptide from the pMHC structure for each epitope, where available (Table 1). Epitopes that have a relatively featureless surface (low SASA), or which are bulged out of the MHC (high SASA) tend to generate a less diverse TCR repertoires (Turner et al., 2005; Turner et al., 2006). Therefore, we expect the most MI to be associated with epitopes that have very high or very low SASA. To test for this, we determined whether a relationship between SASA and MI exists by fitting a parabola with ordinary least squares (Figure 7B). No significant relationship was detected, but structures are available for only 13 out of 22 studied epitopes, which limits the statistical power of the analysis.

### Using co-evolution methods for TCR*αβ* pairing

The correlations of sequence similarity and MI across chains (Figures 4 and 5) resemble those of co-evolving protein families (De Juan et al., 2013). We therefore wondered whether we could use methods developed for co-evolving proteins to select the most likely TCR*β* partner for each TCR*α* within epitope-specific repertoires, when the pairing information is withheld. We adapted two methods: a MI-based method (MI-IPA, Bitbol, 2018), which exploits the MI between co-evolving residue pairs (as in Figure 6), and a graph alignment method (GA, Bradde et al., 2010), which exploits the similarity in phylogenetic trees (in our use-case, similar similarity graphs, as in Figure S13), as well as a combination of the two (GA+MI-IPA, Gandarilla-Pérez et al., 2023).

We optimised the MI-IPA and GA by comparing performance on two different epitopes, GLCTLVAML (EBV) and YLQPRTFLL (SARS-CoV-2, Figures S5 and S6, respectively). We then ran the optimised models on all epitopes. To reduce computational time, we subsampled epitopes with *>* 1000 TCRs to 5 subsamples of 700.

First, we ran the MI-IPA on each epitope-specific repertoire. The results are summarised in Figure 8. In 14/22 epitopes, the MI-IPA performs significantly better than random guessing (we consider it successful for epitopes GILGFVTL and RAKFKQLL but not AVFDRKSDAK based on whether it is significant in at least 3/5 subsamples). As the model was run 10 times, we explored whether the stability of the assignments between repeats could provide a confidence score for each pair. For most epitopes, the majority of pairs are not assigned stably across repeats (Figure S18A), but a few *αβ* are always paired together. Epitopes such as ATDALMFTGF and RLRAEQVK, which show no signal of learning (Figure 8), also seem to have unstable assignments, whilst epitopes such as ASNENMETM and SSPPMFRV, for which the MI-IPA can make predictions, show a large proportion of stable pairs. Notably, stable pairs are enriched for correct pairs for all epitopes tested (Figure S18B). Thus, stability may be useful to assess the reliability of the pairing assignment algorithm (as in Bitbol et al., 2016).

**Figure 8.**
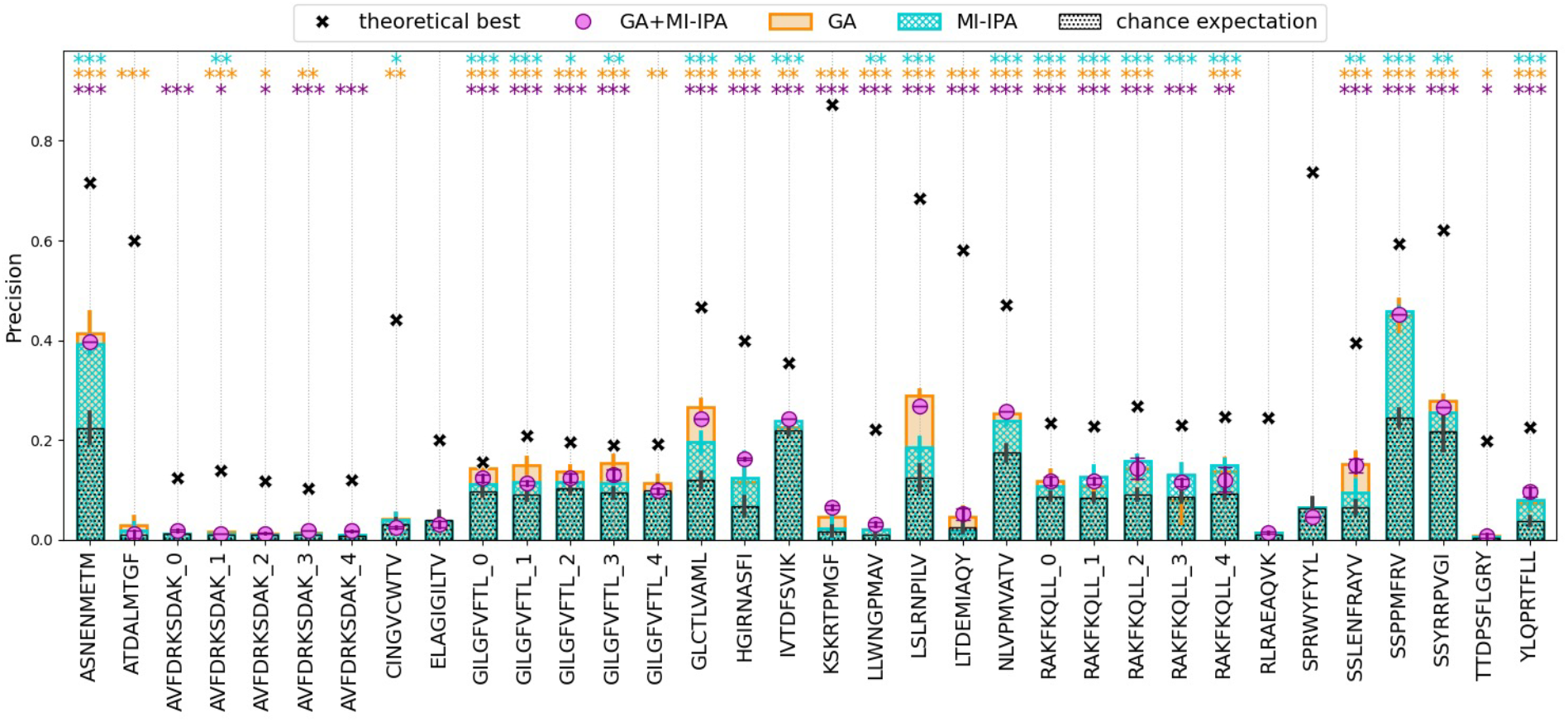
Co-evolution based models perform better than random at pairing TCR*αβ* in epitope-specific repertoires. Precision is calculated over 10 repeats of the MI-IPA or GA+MI-IPA (cyan and magenta, respectively) or 100 repeats of the GA (orange) for all epitopes. For epitopes with *>* 1000 sequences, 5 subsamples of 700 sequences are shown. The IPA is run both with *λ* = 1 (chance expectation, black bars) and *λ* = 0.6 (cyan). Error bars correspond to the standard deviation over the repeats. The black crosses correspond to the theoretical best performance, i.e. the MI-IPA initialised on all correct pairs and run once to pair all sequences. One-sided Student’s t-test is calculated for each epitope for the MI-IPA, GA or GA+MI-IPA using the alternative hypothesis that the model performs better than chance expectation. P-values are indicated by asterisks in the corresponding colour: \*\*\**<* 0.001; 0.001 *≤*\*\**<* 0.01; 0.01 *≤*\**<* 0.05.

We then ran the GA algorithm to pair TCR*αβ* from all available epitope repertoires (Figure 8). Overall, the GA shows significantly better performance than chance expectation in 19/22 epitopes. As for the MI-IPA, epitopes that perform well have more stable assignments than epitopes for which the GA cannot achieve good performance (Figure S19A). Moreover, stable pairs are enriched for correct pairs (Figure S19B).

Finally, we combined the two methods as suggested in Gandarilla-Pérez et al., 2023. Briefly, stable pairs from the GA assignments provide the training set for the MI-IPA. We opted to retain these sequences in the testing classification task, so as to allow the MI-IPA to correct any mistakes in the GA classification (Figure S7). The results for the GA+MI-IPA are shown in Figure 8. The combination of the two methods occasionally marginally improved on either method alone. However, unlike the stability plots for MI-IPA and GA, the GA+MI-IPA shows a larger proportion of stable pairs (Figure S20). We attribute this to the use of a training set in the MI-IPA when combined to the GA as the trajectory of the MI-IPA is less random when a training set is provided. Unfortunately, this has repercussions on the ability to use stability to enrich for correct pairs: for epitopes such as ASNENMETM, GLCTLVAML or LSLRNPILV the precision in the most stable pairs is similar to the overall precision (Figure S20B).

Overall, the pairing algorithms showed a modest but significant ability to identify true pairs of *αβ* sequences. To understand whether the limitation was in the learning step (i.e. the way the algorithm selects pairs does not allow it to find the optimal solution) or in the data (i.e. the signal in the data is too low to achieve pairing), we calculated the theoretical best performance for the MI-IPA. We initialised the MI-IPA with all correctly-paired *αβ* for each epitope. This allows the MI-IPA to immediately see all the rules that govern the pairing at once (the complete set of statistical dependencies). We then ran a single iteration of the MI-IPA to pair all the available *α*/*β* from that epitope repertoire again. We could thus assess whether the MI-IPA correctly pairs all available sequences knowing the pairing rules *a priori* (Figure 8). In all epitopes, the pairing algorithms under-perform compared to the theoretical limit, suggesting that the learning process is not able to extrapolate all the rules that can be learnt from these pairs, i.e. it converges to a different solution. However, the theoretical maximum is well below 1 for all epitope repertoires, suggesting that the signal available from the data is limited, and can recapitulate only some of the rules of TCR*αβ* pairing. We posit that dataset size and difficulty in aligning these sequences may be some of the limiting factors in this approach.

Finally, we wondered if TCR sequence embedding methods or models trained to predict TCR-epitope binding could capture the pairing signal. We therefore predicted chain pairing within an epitope-specific repertoire using TULIP (Meynard-Piganeau et al., 2024), a method developed for binding prediction, and SCEPTR (Nagano et al., 2025), a method for TCR sequence embedding. We find that co-evolution-based models consistently perform better than both TULIP and SCEPTR for the pairing problem (Figure S21), suggesting that they can capture a signal that is often overlooked in other specificity-prediction methods. However, both TULIP and SCEPTR seem to be able to learn some of the pairing signal in their training, as both perform marginally better than chance expectation for a few epitopes.

### Correlates of pairing performance and repertoire characteristics

We sought to understand which factors drive the large range in precision observed in Figure 8 for each model. The MI-IPA is a data-thirsty method, and the total size of the repertoire, as well as the size of the individual repertoires can highly influence performance (Bitbol, 2018), as larger individual repertoire sizes will lead to more combinatorial pairs to be evaluated. Moreover, the final result might be influenced by the MI available in each epitope-specific repertoire (defined as in Figure 5). We therefore correlated MI-IPA performance (calculated on the consensus assignment over 10 repeats, i.e. by taking the modal *β* for each *α*) with each of these factors (Figure 9A). The theoretical maximal, but not the actual performance was inversely correlated to individual repertoire size and total epitope repertoire size, and positively correlated to MI. These 3 variables together can explain over 50% of the variance observed across epitopes in the theoretical best performance, and almost 30% in the learning scenario (multivariate linear regression, Table S2).

**Figure 9.**
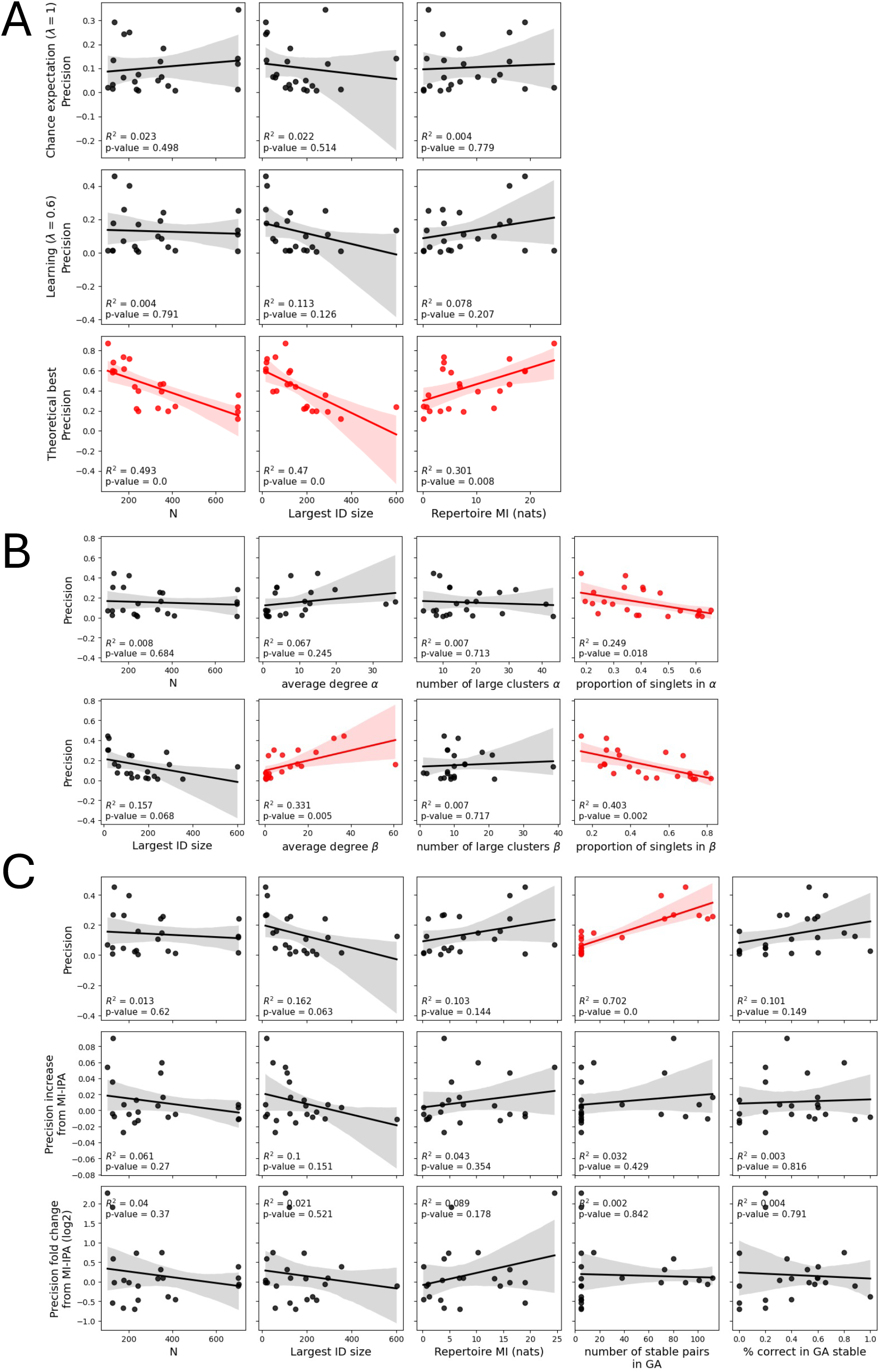
Correlation of model performance and repertoire characteristics. **A)** A linear regression was calculated to evaluate the effect of repertoire size (N), largest individual repertoire size (Largest ID size) and total mutual information (Repertoire MI, calculated between CDR3*α* and CDR3*β* as in Figure 5, without fold change from background) on the precision for MI-IPA (no learning: [confidence = none; *λ* = 1.0; *θ* = 0.6]; learning: [confidence = none; *λ* = 0.6; *θ* = 0.6]; theoretical best: [confidence = none; *λ* = 0.6; *θ* = 0.6; correct pairs as training set]). Subsamples from large repertoires are included as an average, and the repertoire MI is calculated on the complete sample. **B)** A linear regression was calculated to evaluate the effect of N, Largest ID size and similarity network characteristics on the precision for GA. The graph characteristics measured are: average degree of nodes, number of clusters of size *≥* 3 and proportion of sequences that do not have any neighbours (singlets). These are calculated on the graph built by drawing edges between sequences at Levenshtein distance *<* 3 on CDR3*α* and CDR3*β* separately. Subsamples from large repertoires are included as an average, and the graph properties are calculated on each subsample separately. **C)** A linear regression was calculated to evaluate the effect of N, Largest ID size, Repertoire MI, number of stable GA pairs and proportion of the GA training set that is correct (% correct in GA stable) on the precision for GA+MI-IPA. Subsamples from large repertoires are included as an average, the repertoire MI is calculated on the complete sample and the GA stability results are calculated for each subsample separately and averaged. In each panel, the precision is calculated using the modal *β* for each *α* over multiple iterations of the model. Since multiple CDR3*β* may be selected the same number of times for one CDR3*α*, one is selected at random. To mitigate for variation, the mode selection is run 100 times and the average precision is used. In each panel, the *R*^2^ for the regression is shown and the scatterplots are red when p-value for the F-statistics of the regression is *<* 0.05. The solid line shows the calculated regression and the shaded area the 95% confidence interval of the prediction.

To understand which factors influence GA performance, we correlated its performance (calculated on the consensus assignment over 100 repeats) with repertoire size, largest individual size, as well as characteristics describing the sequence similarity networks that can be generated from each of these epitopes (calculated using pairwise Levenshtein distance, threshold *<* 3). The results are shown in Figure 9B. Interestingly, performance correlates with the properties of the sequence similarity graph: the more sequence clustering is observed (higher average degree, smaller proportion of singlets), the better the GA is at predicting outcome. The model built with these variables can explain almost 80% of the variance between epitopes (Table S3).

Finally, we looked at factors that might impact the performance of the GA+MI-IPA. As for the MI-IPA, we calculated the effect of repertoire size, largest individual repertoire size and repertoire MI. Moreover, we extracted the number of stable pairs from the GA and the percentage of the GA stable set that is correct. We correlated these factors with both the precision, increase in precision from using MI-IPA on its own, as well as the fold change between the precision of the MI-IPA and the precision of the GA+MI-IPA, to see if we could explain why some epitopes benefit from the initial GA step and some do not. Figure 9C shows the results for each variable independently. The number of stable pairs in the GA is significantly correlated with precision of the GA+MI-IPA. Interestingly, none of these factors can explain why certain epitopes improve on the MI-IPA and others do not. When combining these variables in a multivariate regression, they can explain over 80% of the observed variance in precision of the GA+MI-IPA, but they are unable to explain the variance in improvement compared to MI-IPA alone (Table S4).

## Discussion

We report conserved intra- and inter-chain contacts between the TCR CDR loops. Specifically, we observed intra-chain interactions between all CDR loops and inter-chain interactions between the conserved FR2 on each chain, and at the interface between the two CDR3s. These interactions are conserved in TCRs both unbound and bound to class I and class II pMHCs, although the exact topology of the interactions can change upon binding. We did not detect conserved interactions between the germline-encoded loops on the two chains, consistent with the idea that there are no significant constraints in TRAV/TRBV pairing at the repertoire level (Dupic et al., 2019; Shcherbinin et al., 2020). Whilst previous structural studies have observed interactions between TCR*α* and TCR*β* chains, this is, to our knowledge, the first systematic review of all available structures, which shows that these interactions are a conserved feature of the folded TCR structure, and are present in both the bound and unbound state. The TCR antigen-binding site is therefore shaped by interactions between the six CDR loops, and these interactions may play a role in determining the thermodynamics and kinetics of antigen binding.

Having observed that the CDR regions of the two chains are consistently packed in close proximity in the folded TCR structure, we hypothesised that this might enforce specific conformations and thus restrict sequence diversity in the context of sets of TCRs with shared pMHC specificity. We found several strands of evidence supporting this prediction. First, using a measure of pairwise sequence similarity, we find that epitope-specific TCRs with similar *α* sequences have paired *β* chains with restricted sequence diversity (and vice-versa). Moreover, we detected consistent MI between the CDR3*α* and the CDR3*β* sequences within sets of epitope-specific TCRs. Whilst we have taken steps to reduce bias due to batch effects or finite-size effects in the MI estimation, we cannot rule out that these may still partially impact our estimations. The mutual restriction seen between TCR*α* and *β* sequence diversity in the context of shared specificity is reminiscent of the ‘light chain coherence’ described in antibodies (Jaffe et al., 2022). Our results differ from those in Shcherbinin et al., 2020, as they did not find evidence for restriction of TCR chain pairing even within epitope-specific repertoires. This discrepancy can likely be attributed to the inclusion of the CDR3 sequence in our analysis. This suggestion is consistent with the result that the CDR3*α* and CDR3*β* carry synergistic information about epitope binding, greater than the synergy measured between V*α* and V*β* genes (Henderson et al., 2024). The diversity restrictions discussed above were observed across most epitopes tested. However, there was considerable variance in the interaction strength detected when comparing TCRs binding different pMHCs, discussed in more detail below.

We reasoned that the sequence constraints observed on TCR*αβ* pairs are somewhat analogous to constraints found between co-evolving protein families, which share similarity in their phylogenetic trees, and show sequence correlations at the residue level (Korber et al., 1993; Fryxell, 1996; Dunn et al., 2008; De Juan et al., 2013; Ochoa & Pazos, 2014). A number of approaches have been developed to predict pairing between co-evolving interacting protein families, showing that such a signal is learnable. We thus wondered whether the TCR*αβ* pairing signal can be learnt, as this would have practical applications in predicting chain pairing from TCR*α* and TCR*β* chains sequenced independently. To test this, we reframed the TCR*αβ* pairing problem as a co-evolution problem: we consider the multiple TCR*α* and TCR*β* chains which bind to the same epitope as ‘paralogs’ from interacting protein families, while similar TCR*α* (or *β*) sequences that bind the same epitope but are found in different individuals are ‘orthologs’. We adapted three co-evolution based methods to try and predict TCR*αβ* pairing (Bradde et al., 2010; Bitbol, 2018; Gandarilla-Pérez et al., 2023) and applied them in the simplest possible scenario: by using lists of TCR*α* and TCR*β* previously annotated for a single epitope. Except for the theoretical best scenario, a training set of know TCR*αβ* pairs was never provided to the pairing algorithms, so that we could evaluate each model’s ability to make *de novo* pairing predictions. We were able to detect a significant improvement in pairing performance over random assignment in all epitopes we tested except three (ELAGIGILTV, RLRAEAQVK and SPRWYFYYL) using at least one of the three methods. Notably, previous studies have suggested that the TCR recognition of epitope ELAGIGILTV is dominated by the TCR*α* chain (Trautmann et al., 2002; Dietrich et al., 2003), and may be largely independent of TCR*β*.

Overall, the co-evolution based methods show better performance in the pairing task than existing models for epitope prediction (Meynard-Piganeau et al., 2024) or TCR sequence embedding (Nagano et al., 2025), suggesting they are learning an alternative signal, which may be important for prediction. However, the classification performance remains poor, precluding current use of the pairing algorithm for practical applications. The addition of more pairs for each epitope will likely increase performance, as these methods tend to be data-thirsty (Bitbol, 2018). The presented methods have a fairly long run time and we have therefore not benchmarked them on larger sets. Faster methods have been developed (see for instance, Lupo et al., 2025) and may work better on larger datasets. Technical improvements might also be possible to achieve better pairing. For example, in some cases the addition of GA did not improve, or even worsened, the performance of the MI-IPA. We speculate that this may occur if the GA places the MI-IPA initial cache at a local MI maximum, which the MI-IPA cannot escape and thus improve upon. Future studies could explore better ways to use the results from the GA to initialise the MI-IPA. On the other hand, it might be possible to boost performance by including information other than TCR sequence, such as clonotype abundance, as we expect chains that come from the same clone to have similar abundance in the same sample, and vary across samples in a correlated manner. Finally, we assume all the unobserved pairs in the data to be non-binders. However, binding of predicted but unobserved pairs was never tested experimentally, and we may thus hypothesise that the pairing methods are indeed identifying new (and maybe optimised) binding pairs.

Importantly, however, we do not think that lack of data or technical improvements of the algorithms are the only limiting factors in achieving successful pairing. The algorithms we use match chains based only on maximizing mutual information or graph-based phylogenetic similarity, but the interchain interaction is only one factor in determining the selection of *α* and *β* chains, and their pairing. Each chain independently interacts with the pMHC, and is therefore subject to independent selection forces by pMHC, which may in some cases be stronger than the forces imposed by pairing. In the co-evolutionary context, interacting proteins evolve over time, and thus existing examples presumably represent optimal (or optimised) solutions to the interaction. In the TCR context, paired *αβ* do not co-evolve, but rather get selected if they overcome a threshold of functionality. As such, these pairs may not represent the optimal solution to the interaction, and instead they may be a functional (and likely sub-optimal) solution that was selected. Indeed, our set includes repeated sequences in both the CDR3*β* and CDR3*α*, which suggests that multiple functional solutions are possible for the same chain (Table S1). We propose that the extent to which the *α* and *β* chains are interdependent, and the extent to which they can contribute independently to binding energy, varies widely between epitopes. However, we posit that, for many epitopes, interchain dependencies captured by MI or GA, are a (underappreciated) component of the overall TCR/pMHC binding.

Different epitopes show different levels of MI, different patterns of where most of the MI is contained, and different performance of the pairing algorithms. We hypothesise that this variation may be explained by both intrinsic and extrinsic features. Intrinsic features reflect physical properties of the TCR/pMHC complex (e.g. more or less contacts, binding constraints etc.). For example, the feature-ness of a peptide can impact the diversity of the associated repertoire (Turner et al., 2005; Turner et al., 2006), although we were unable to detect any correlation between feature-ness and MI. Extrinsic features include size of the epitope-specific repertoires, as well as sequence publicity and similarity within a repertoire. We found significant correlations between the effective set size and the MI of the repertoire: the smaller the effective set size, the higher the MI. We speculate this is due to the presence of fewer, larger clusters, which allow us to better recapitulate the binding rules for each in the MI. This is consistent with the positive correlation between the average degree of the CDR3 similarity networks and the performance of the GA algorithm. Overall, we still do not have a full understanding of the factors which determine the relationship between TCR*α* and *β* interaction and pMHC. A much larger selection of pMHC than the 22 examples available and studied here will need to be analysed to answer these questions.

In conclusion, we identify extensive, and in some cases highly conserved interactions between the CDR sequences which form the binding surface of a TCR, and which we predict will affect the kinetics and thermodynamics of binding. The interactions may reduce the entropy of the TCR prior to binding, and thus increase TCR affinity, on-rate and specificity at the expense of reduced breadth. Further experiments to measure the dynamics of TCR, before and after binding, will be required to validate these predictions, which have important implications in the context of the design of TCR-based therapeutics. In addition, we measure broader sequence constraints on TCR*αβ* pairing in the context of antigen specific TCR sets. We provide some initial indications of how these constraints may be used to help to correctly pair *α* and *β* sequences when such pairing is not available experimentally. Understanding the nature of these constraints may also have broader implications for the ultimate goal of predicting TCR/pMHC binding.

## Data availability

No new data was generated with this paper. The data analysed can be downloaded from VDJdb (https://vdjdb.cdr3.net/), the Structural T Cell Receptor Database (STCRDab, https://opig.stats.ox.ac.uk/webapps/stcrdab-stcrpred/) and the Observed TCR Space database (OTS, https://opig.stats.ox.ac.uk/webapps/ots/). Preprocessed data from Tanno et al., 2020 was obtained from the authors.

## Code availability

The code needed to reproduce the analysis in this manuscript is available at https://github.com/mm523/TCRab-pairing.

## Acknowledgements

A version of the figures and results in this work were previously reported in M.M.’s PhD thesis (Milighetti, 2023). The authors acknowledge the use of the UCL Myriad High Performance Computing Facility (Myriad@UCL), and associated support services, in the completion of this work. This work was supported by Cancer Research UK through a Non-Clinical Training Award to M.M. [A29287] and by grants from the Rosetrees Foundation and the UCLH Biomedical Research Centre to B.C.. Y.N. was supported by the Cancer Research UK City of London Centre (grant number BCCG1C8R). U.H. was supported by NIAID program project grant P01 AI106697 and the European Union’s Horizon 2020 Research and Innovation Program under grant agreement 825821 and by Israel Science Foundation (ISF) grant 1327/22. The work of A. T.-M. was supported in parts by the Royal Free Charity. A.-F. B. thanks the European Research Council (ERC) for funding under the European Union’s Horizon 2020 research and innovation programme (grant agreement No. 851173, to A.-F. B.).

## Author contributions

M.M. and B.C. conceived the study. M.M., Y.N., J.H., U.H., A. T.-M., A.-F. B. and B.C. developed the methodology. M.M. wrote the code with contributions from Y.N.. M.M. carried out the analyses with contributions from Y.N., J.H., U.H., A. T.-M., A.-F. B. and B.C.. M.M., Y.N., J.H., U.H., A. T.-M., A.-F. B. and B.C. interpreted the results. M.M. and B.C. wrote the first version of the manuscript, and M.M., Y.N., J.H., U.H., A. T.-M., A.-F. B. and B.C. reviewed and edited it.

## Supplementary material

**Table S1:** Summary of VDJDb set used for the pairing algorithms. For large epitopes that were subsampled for pairing, composition of each subsample is shown.

| Epitope | Number of entries | Unique CDR3α | Unique CDR3β | Number of individuals |
| --- | --- | --- | --- | --- |
| **ASNENMETM** | 201 | 157 | 111 | 28 |
| **ATDALMTGF** | 125 | 99 | 111 | 1 |
| **AVFDRKSDAK** | 1699 | 1552 | 1584 | 7 |
| subsample 0 | 700 | 661 | 670 | 7 |
| subsample 1 | 700 | 663 | 671 | 6 |
| subsample 2 | 700 | 657 | 662 | 5 |
| subsample 3 | 700 | 653 | 679 | 6 |
| subsample 4 | 700 | 657 | 672 | 6 |
| **CINGVCWTV** | 226 | 214 | 218 | 7 |
| **ELAGIGILTV** | 380 | 348 | 370 | 14 |
| **GILGFVFTL** | 1894 | 1082 | 980 | 56 |
| subsample 0 | 700 | 477 | 442 | 48 |
| subsample 1 | 700 | 487 | 460 | 50 |
| subsample 2 | 700 | 488 | 427 | 50 |
| subsample 3 | 700 | 464 | 436 | 50 |
| subsample 4 | 700 | 475 | 423 | 44 |
| **GLCTLVAML** | 345 | 228 | 239 | 20 |
| **HGIRNASFI** | 243 | 187 | 180 | 13 |
| **IVTDFSVIK** | 704 | 526 | 517 | 7 |
| **KSKRTPMGF** | 103 | 85 | 70 | 1 |
| **LLWNGPMAV** | 235 | 202 | 220 | 2 |
| **LSLRNPILV** | 127 | 109 | 111 | 12 |
| **LTDEMIAQY** | 124 | 115 | 119 | 2 |
| **NLVPMVATV** | 357 | 305 | 318 | 48 |
| **RAKFKQLL** | 1200 | 649 | 650 | 3 |
| subsample 0 | 700 | 411 | 407 | 3 |
| subsample 1 | 700 | 417 | 405 | 3 |
| subsample 2 | 700 | 410 | 411 | 3 |
| subsample 3 | 700 | 402 | 420 | 3 |
| subsample 4 | 700 | 419 | 406 | 3 |
| **RLRAEAQVK** | 412 | 389 | 394 | 4 |
| **SPRWYFYYL** | 175 | 169 | 174 | 10 |
| **SSLENFRAYV** | 350 | 240 | 286 | 19 |
| **SSPPMFRV** | 133 | 100 | 58 | 14 |
| **SSYRRPVGI** | 177 | 148 | 153 | 32 |
| **TTDPSFLGRY** | 242 | 231 | 232 | 1 |
| **YLQPRTFLL** | 333 | 284 | 304 | 10 |

**Table S2:** Factors affecting MI-IPA performance. A multivariate linear regression was calculated to evaluate the effect repertoire size (N), largest individual (Largest ID size) and total mutual information (Repertoire MI, calculated as in Figure 5 between CDR3*α* and CDR3*β*) on the precision. Subsamples from large repertoires are included as an average, and the repertoire MI is calculated on the complete sample. Each independent variable was normalised by subtracting the mean and dividing by the mean to make the derived coefficients more comparable 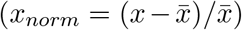. The *R*^2^ and adjusted 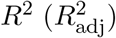 for the regression are shown, as well as the F-score p-value. P-values associated with each coefficient are shown as asterisks: \*\*\**<* 0.001; 0.001 *≤*\*\**<* 0.01; 0.01 *≤*\**<* 0.05.

| | N | Largest ID size | Repertoire MI (nats) | $R^2$ | $R^2_{\text{adj}}$ | F-score p-value |
| --- | --- | --- | --- | --- | --- | --- |
| **No-learning background:**<br>confidence = none; $\lambda = 1.0$; $\theta = 0.6$ | 0.1243<br>(*) | -0.0854 | 0.0118 | 0.230 | 0.102 | 0.185 |
| **Learning model:**<br>confidence = none; $\lambda = 0.6$; $\theta = 0.6$ | 0.1327 | -0.1116<br>(*) | 0.0366 | 0.276 | 0.155 | 0.113 |
| **Theoretical best:**<br>confidence = none; $\lambda = 0.6$; $\theta = 0.6$ | -0.1219 | -0.0731 | 0.0567 | 0.574 | 0.502 | 0.00129 |

**Table S3:** Factors affecting GA performance. A multivariate linear regression was calculated to evaluate the effect repertoire size (N), largest individual (Largest ID size) and similarity network characteristics on the precision for each model. The graph characteristics measured are: average degree of nodes, number of clusters of size *≥* 3 and proportion of sequences that do not have any neighbours (singlets). These are calculated on the graph built by drawing edges between sequences at Levenshtein distance *≤* 3 on both CDR3*α* and CDR3*β*. Epitopes for which 5 subsamples are available are averaged across subsamples, and each of the metrics is calculated on the specific subsample. Each independent variable was normalised by subtracting the mean and dividing by the mean to make the derived coefficients more comparable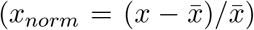. The *R*^2^ and adjusted 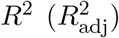 for the regression are shown, as well as the F-score p-value. P-values associated with each coefficient are shown as asterisks: \*\*\**<* 0.001; 0.001 *≤*\*\**<* 0.01; 0.01 *≤*\**<* 0.05.

| | N | Largest ID size | Average degree of node | Number of large clusters | Proportion of singlets | $R^2$ | $R^2_{\text{adj}}$ | F-score p-value |
| --- | --- | --- | --- | --- | --- | --- | --- | --- |
| **GA:**<br>distance=lev; $k=20$ | -0.3507<br>(*) | -0.1282 | $\alpha$: 0.0608<br>$\beta$: 0.1062 (**) | $\alpha$: 0.3761<br>$\beta$: 0.0414 | $\alpha$: 0.2471<br>$\beta$: -0.0251 | 0.799 | 0.675 | 0.002 |

**Table S4:** Factors affecting GA+MI-IPA performance. A multivariate linear regression was calculated to evaluate the effect repertoire size (N), largest individual (Largest ID size), total mutual information (Repertoire MI, calculated as in Figure 5 between CDR3*α* and CDR3*β*), number of stable GA pairs and proportion of the GA training set that is correct (% correct in GA stable) on the precision. Each independent variable was normalised by subtracting the mean and dividing by the mean to make the derived coefficients more comparable 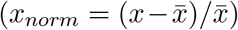 The *R*^2^ and adjusted 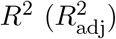 for the regression are shown, as well as the F-score p-value. P-values associated with each coefficient are shown as asterisks: \*\*\**<* 0.001; 0.001 *≤*\*\**<* 0.01; 0.01 *≤*\**<* 0.05.

| | N | Largest ID size | Repertoire MI (nats) | Number of stable pairs in GA | % correct in GA stable | $R^2$ | $R^2_{\text{adj}}$ | F-score p-value |
| --- | --- | --- | --- | --- | --- | --- | --- | --- |
| **GA+MI-IPA model precision:**<br>distance=lev; $k=20$;<br>GA-thresh=0.95; re-pair=True | -0.0272 | -0.0039 | 0.0418<br>(*) | 0.0814<br>(***) | 0.0502 | 0.828 | 0.774 | $1.29 \times 10^{-5}$ |
| **precision increase from MI-IPA:**<br>precision of GA+MI-IPA<br>–<br>precision of MI-IPA | -0.0088 | -0.0069 | -0.0024 | 0.0012 | 0.0098 | 0.147 | -0.119 | 0.734 |
| **precision ratio between GA+MI-IPA and MI-IPA:**<br>$\log_2(\frac{\text{precision of GA+MI-IPA}}{\text{precision of MI-IPA}})$ | -0.2301 | 0.0804 | 0.2342 | -0.0312 | 0.1054 | 0.103 | -0.177 | 0.863 |

**Figure S1:**
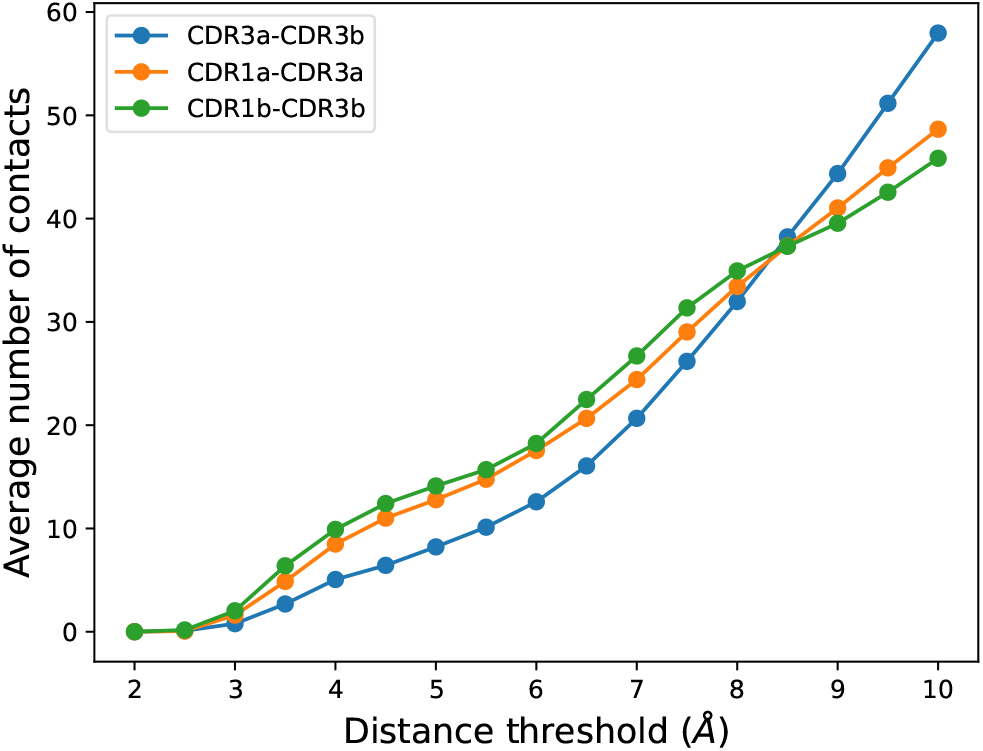
Effect of distance threshold selection on the average number of contacts. Number of observed contacts between CDRs are calculated for each of 340 unique PDB structures at different threshold.

**Figure S2:**
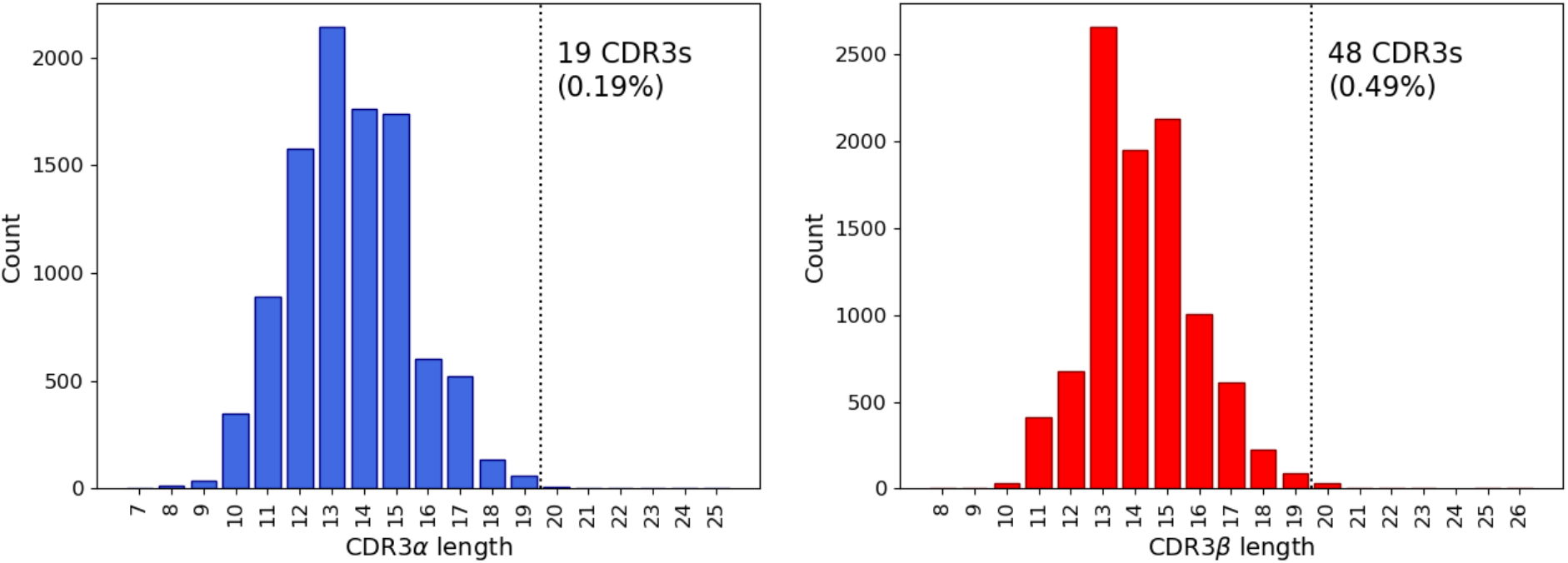
Length of CDR3*α* and CDR3*β* sequences in VDJDb dataset. Distribution of CDR3 lengths (without padding) in the *α* (left) and *β* (right) chains. The dotted line shows the length threshold chosen as maximum allowed length, and the text indicates how many CDR3s were removed and what percentage they correspond to.

**Figure S3:**
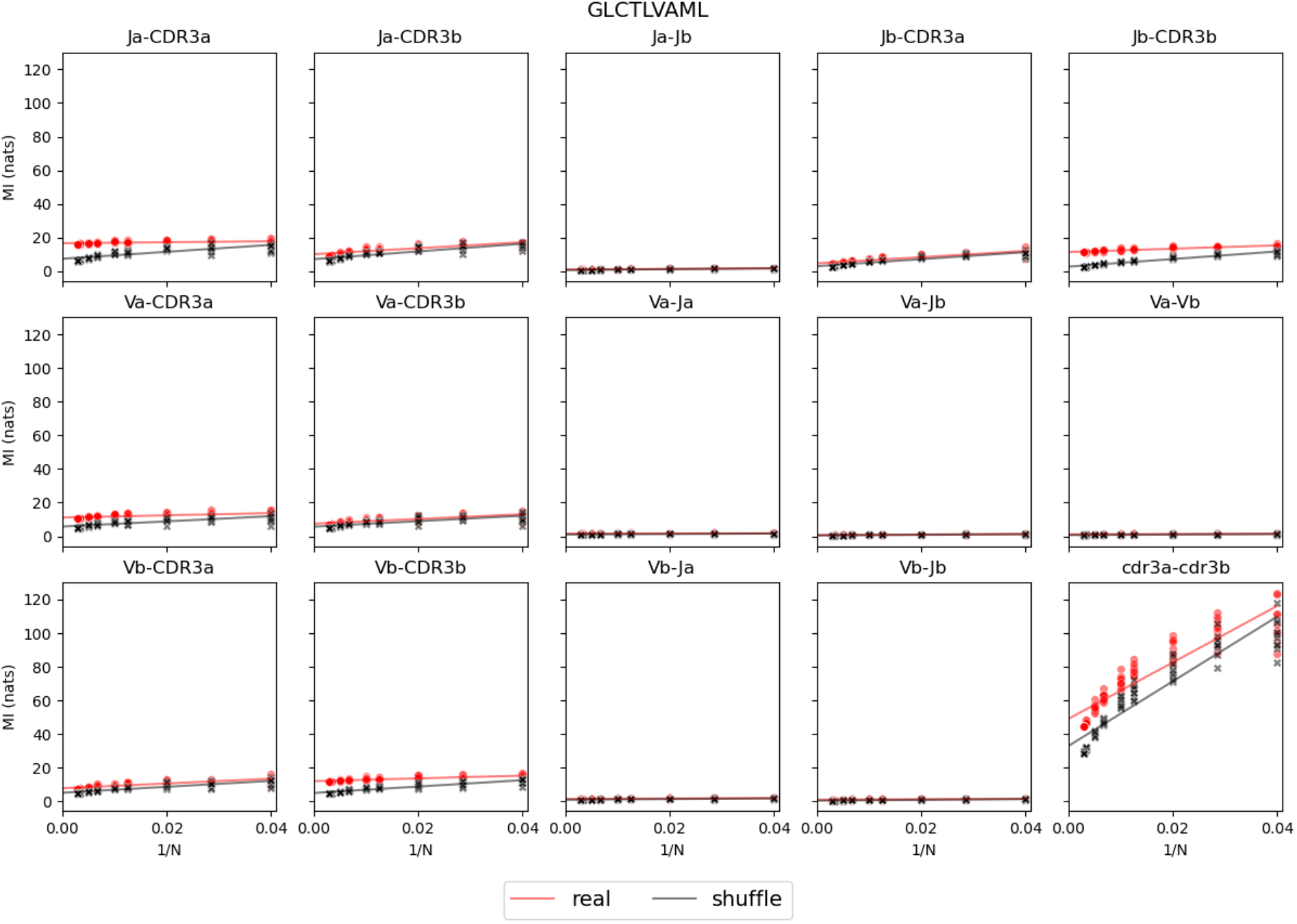
Estimation of mutual information for epitope GLCTLVAML. For each pair of TCR components (indicated in the title of each plot), mutual information (MI) was calculated at multiple subsamples for both the real set (red) and a shuffle (black). Each subsample was repeated 10 times. A line was fit through the point and MI was estimated as the y-intercept for both real set and shuffle.

**Figure S4:**
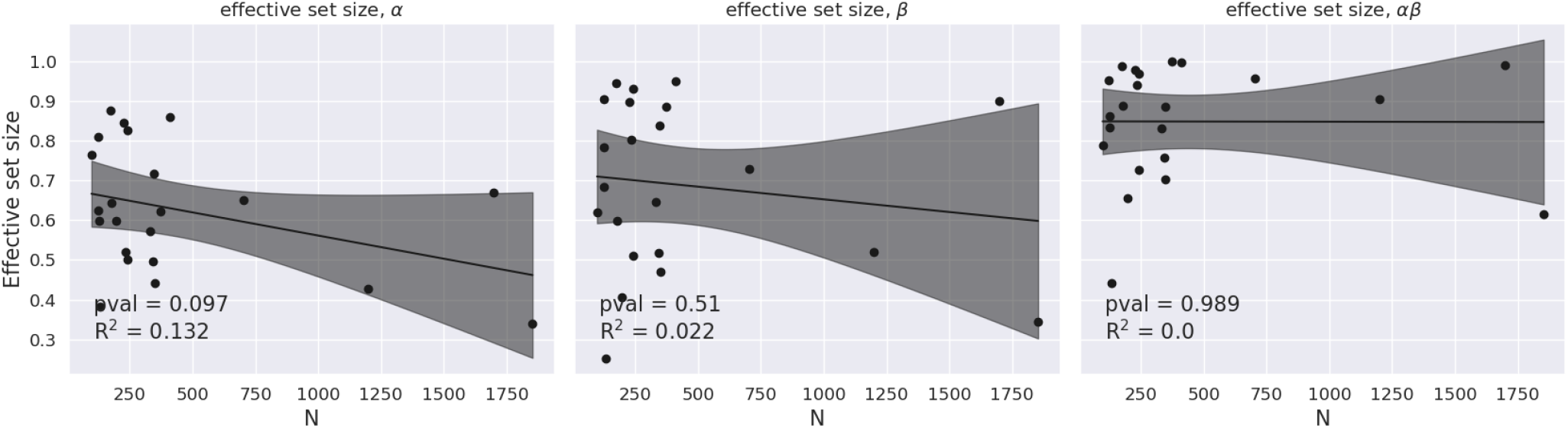
Correlation of repertoire size and effective set size. For each epitope, CDR3*α*, CDR3*β* and CDR3*αβ* effective set size is correlated with total repertoire size (*N*). Effective set size was calculated by grouping all sequences with Hamming distance of *<* 2 on each chain or *<* 3 for the paired chains, normalised by *N* . Correlation was calculated by fitting a linear regression. p-value for the F-test and *R*^2^ was calculated for each fit. The shaded area represents the 95% CI of the fit.

**Figure S5:**
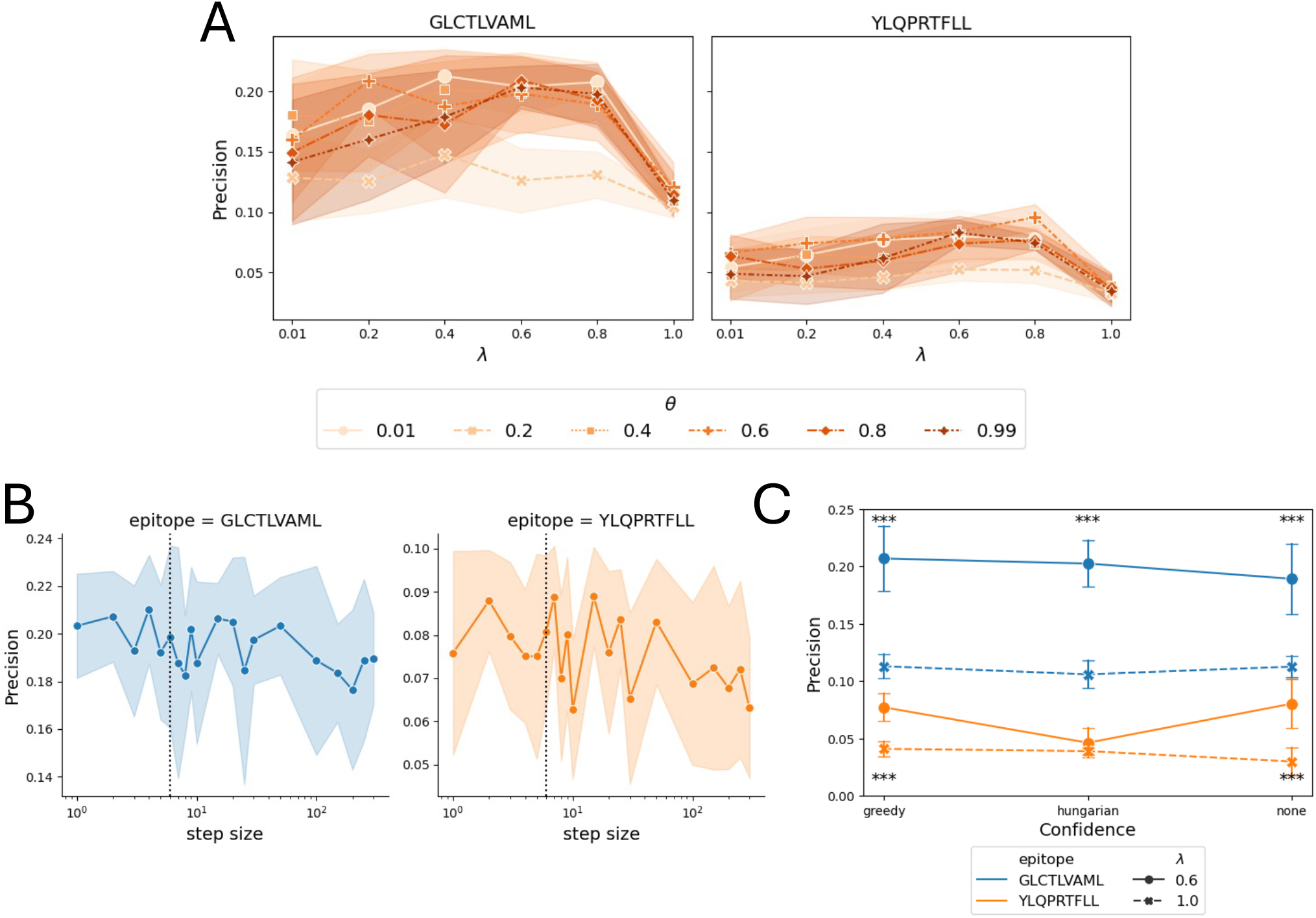
Optimisation of parameters for MI-IPA. All parameters are as in Bitbol, 2018, except that we add a scenario where no ‘confidence’ is calculated. This is a greedy assignment (as in “greedy”), but no confidence score is calculated for the pairing. Rather, the scoring for each pairing is used raw to rank the pairs. Shaded areas or error bars represent the standard deviation around the mean (solid line) for 10 repeats of each model. **A)**: *θ* and *λ* (controlling sampling correction and effective set size calculation, respectively); **B)**: step size; **C)**: confidence calculation algorithms. One-sided t-tests are calculated to compare performance of each model to its no-learning scenario (achieved by setting *λ* = 1). pvalues: \*\*\**<* 0.001; 0.001 *≤*\*\**<* 0.01; 0.01 *≤*\**<* 0.05.

**Figure S6:**
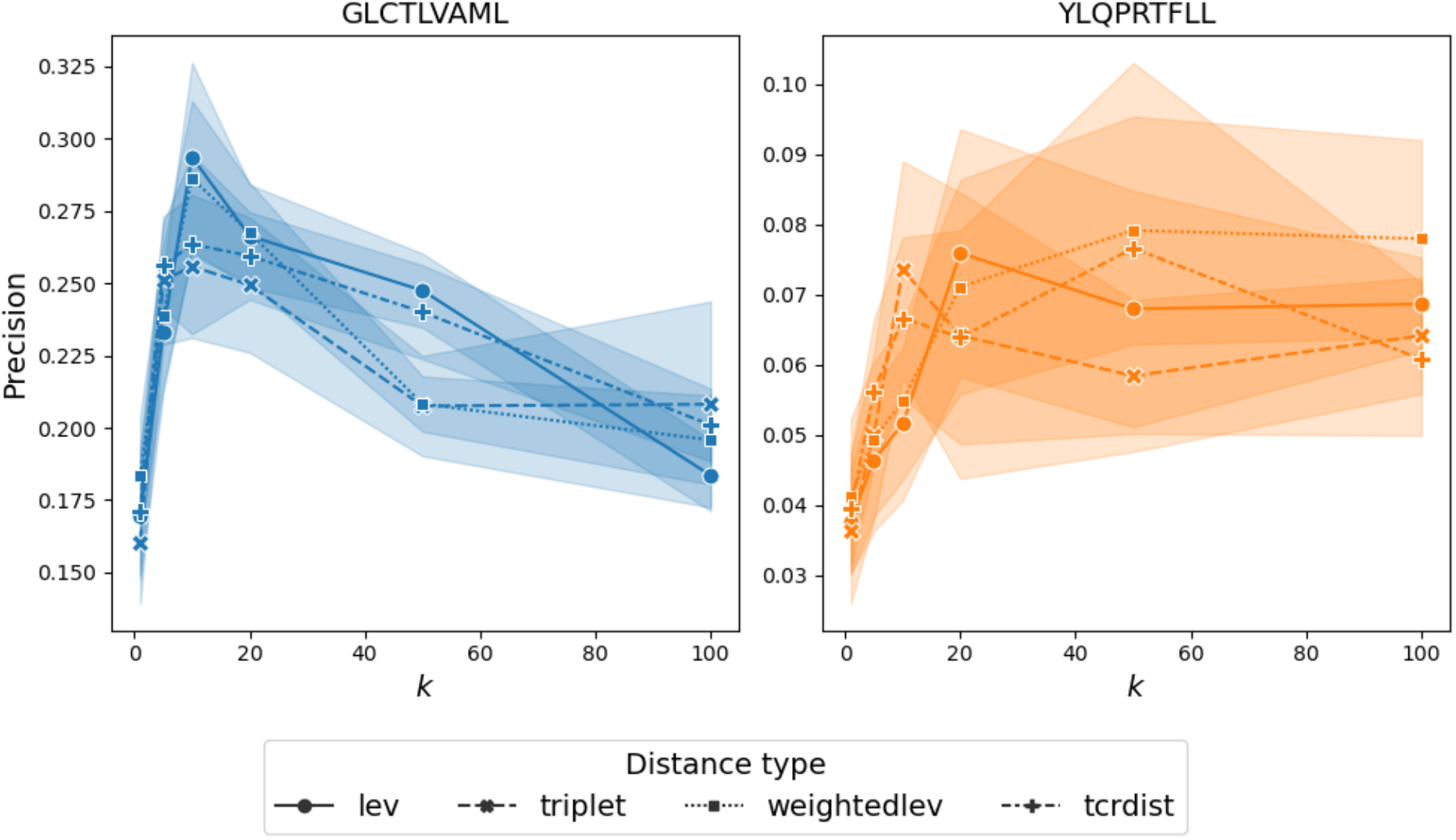
Impact of *k* and distance function choice on GA pairing. The GA was run with a range of *k* and 4 different distance metrics to evaluate their impact on the two benchmarking epitopes. Precision was calculated for each setting. The shaded areas show the standard deviation around the mean (solid line) for 100 repeats of the model.

**Figure S7:**
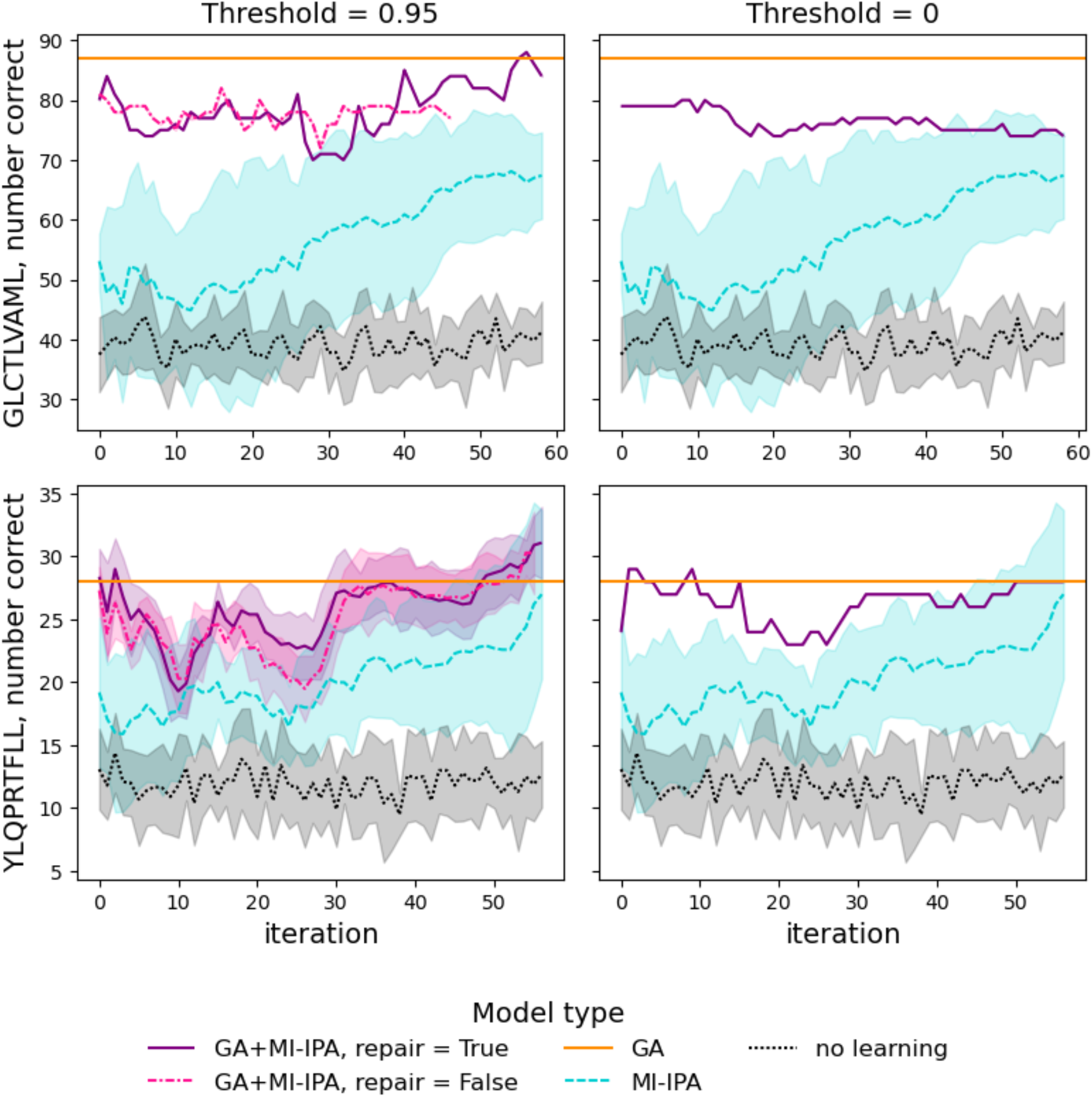
Learning of the GA+MI-IPA over iterations with different settings. Performance of combining the GA (Levenshtein distance, *k* = 20) with MI-IPA (*θ* = 0.6, no confidence, step size = 6) is shown for the two benchmarking epitopes. The orange horizontal line shows the number of correct pairs in the GA consensus assignment which was used to select the training sets. The GA assignment is used as training for the MI-IPA, either by taking the complete assignment (threshold = 0, right column), or only the stable pairs (threshold = 0.95, left column). The MI-IPA is then allowed to pair all remaining *β* to all remaining *α* (pink dotted line with fewer iterations) or pair all *α* and *β*, including the ones in the training set (purple line). Performance of the MI-IPA on its own is also shown, with *λ* = 0.6 (cyan line) or *λ* = 1 (no learning, black dotted line). Shaded areas show the standard deviation around the mean (solid line).

**Figure S8:**
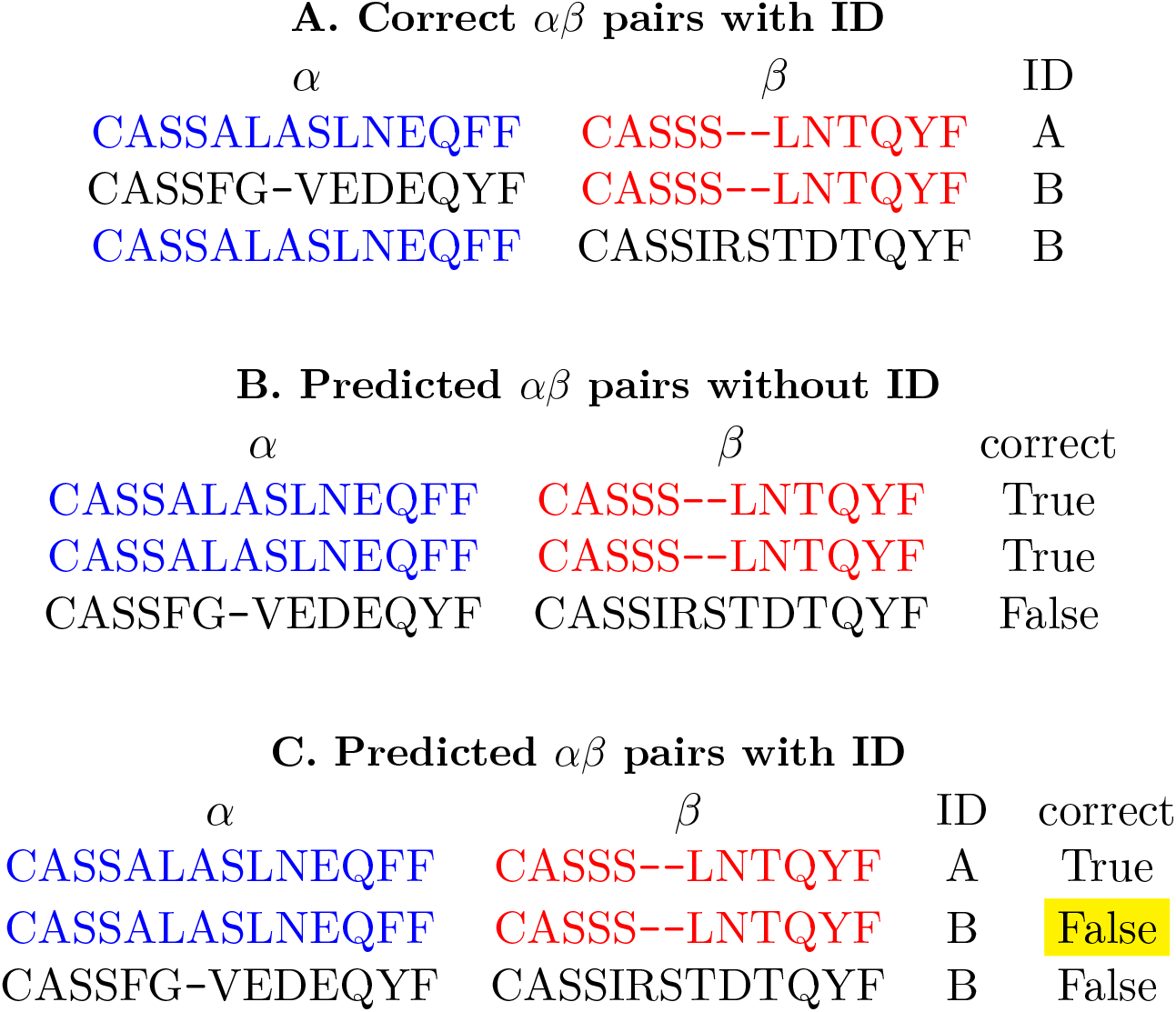
Example assignment with repeat sequences and ID. **A)** Correct pairs to be found. The blue *α* appears twice in the two individuals, but paired with two different *β*. The red *β* also appears twice, once paired with the blue *α*. **B)** Pairs assigned by the algorithm, disregarding individual information. Here, the blue *α*/red *β* pair appears twice. In the precision calculation, this will be counted twice as correct, giving a precision of 0.66. **C)** Pairs assigned by the algorithm, including individual information. Here, the blue *α*/red *β* pair appears twice. Because the blue *α*/red *β* pair does not appear in individual B, the pair is considered correct only in individual A.

**Figure S9:**
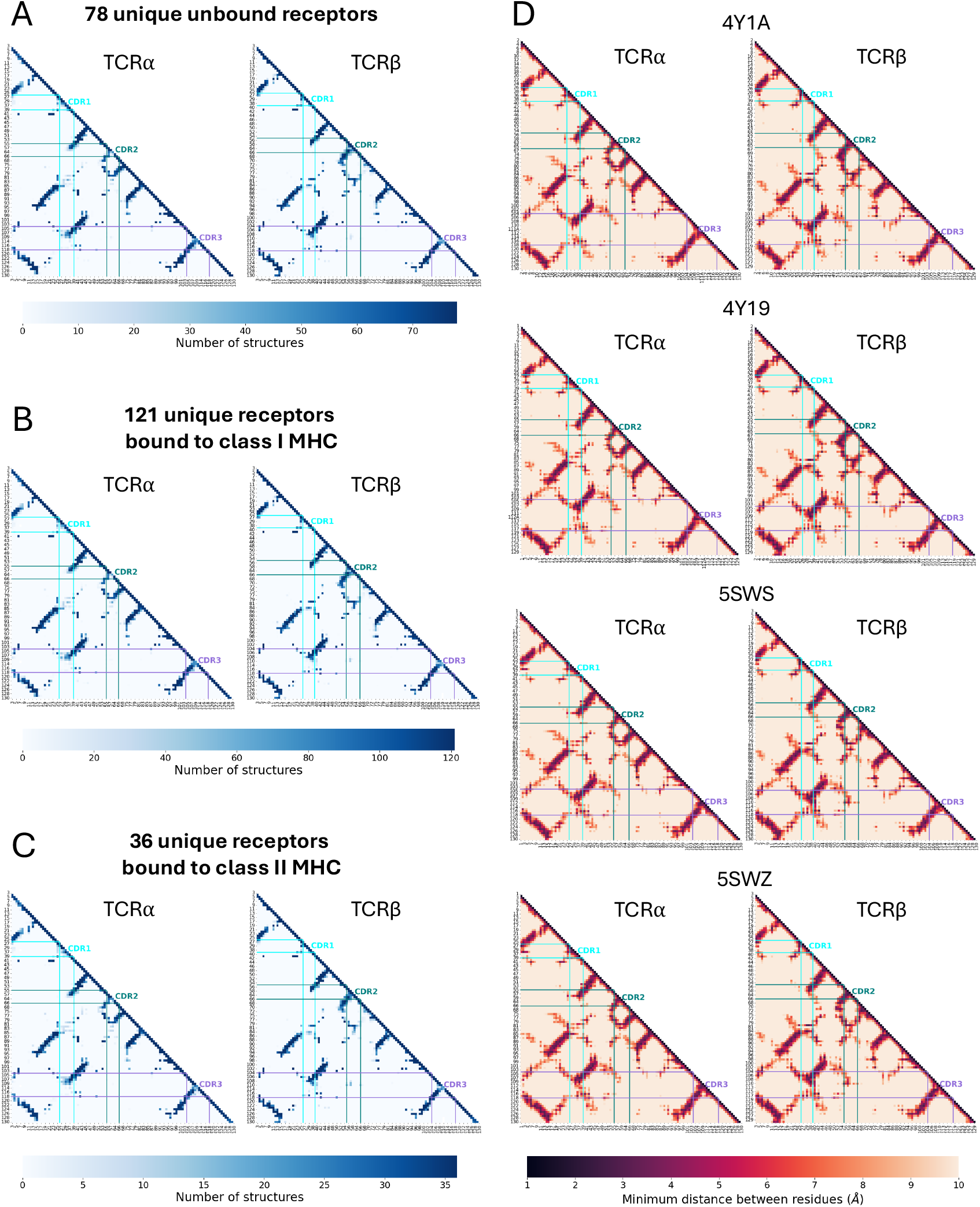
Intra-chain contacts of unbound TCRs and TCRs bound to class I and class II MHC. Intra-chain contact maps were calculated for the TCR *α* and *β* chain of 78 unique unbound TCRs **(A)** and 157 unique bound TCRs to either class I **(B)** or class II MHC **(C)**. A contact was defined as minimum distance between residues *<* 5Å. The heatmap shows the number of structures that are found to make contact at each pair of positions. Only positions present in *>* 75% of structures according to the IMGT numbering scheme are shown. Only the bottom half of each distance matrix is shown as it is symmetrical across the diagonal. In each heatmap, the start and end positions of CDR1, 2 and 3 are highlighted. **D)** Intra-chain contact maps for the TCR *α* and *β* chain for 4 TCRs known to bind MHC with reversed orientation. The distance (Å, colour bar) is calculated between all pairs of atoms in the two residues, and the minimum distance is shown.

**Figure S10:**
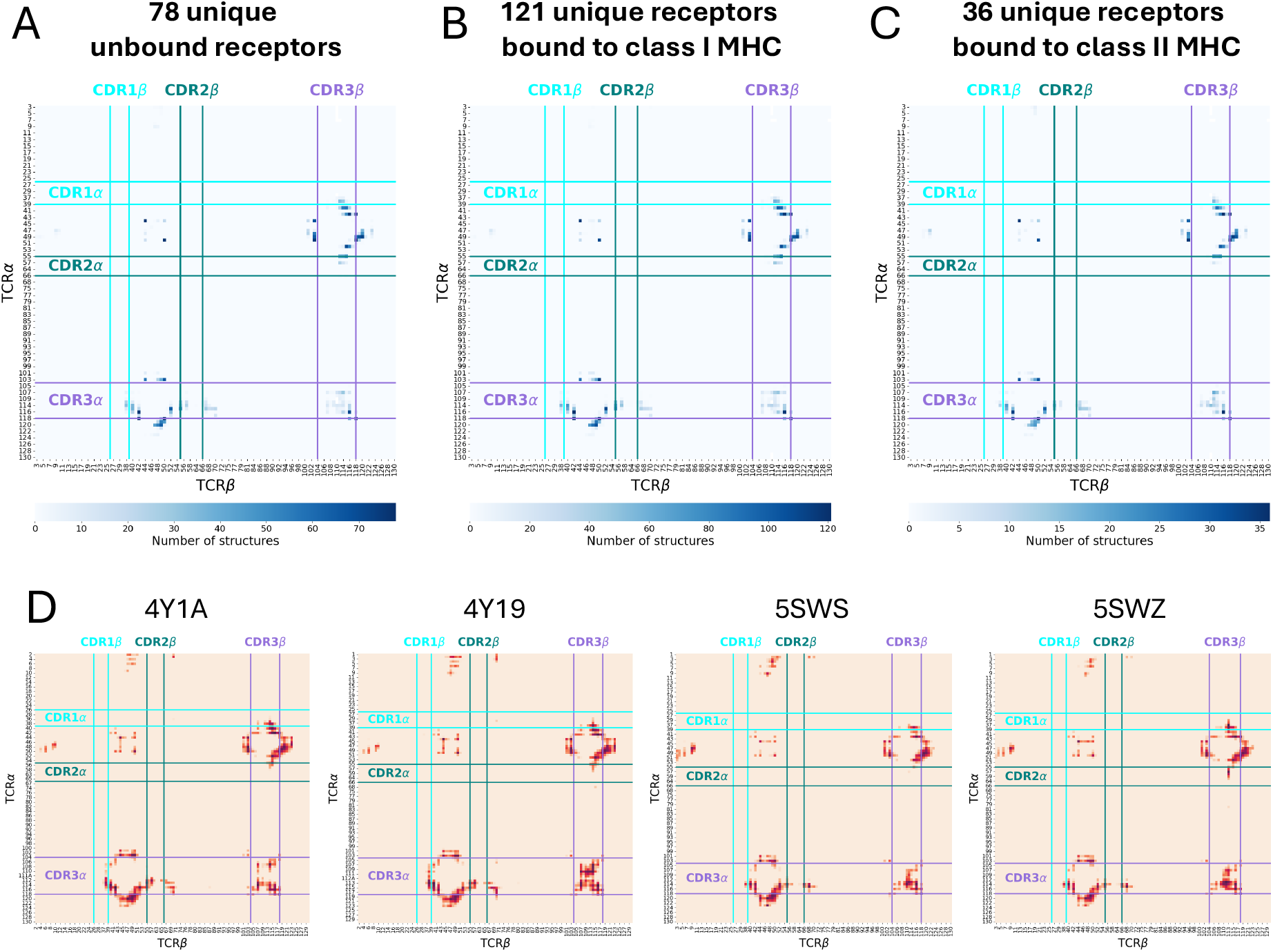
Inter-chain contacts of unbound TCRs and TCRs bound to class I and class II MHC. Inter-chain contact maps were calculated for 78 unique unbound TCRs **(A)** and 157 unique bound TCRs to either class I **(B)** or class II MHC **(C)**. A contact was defined as minimum distance between residues *<* 5Å. The heatmap shows the number of structures that are found to make contact at each pair of positions. Only positions present in *>* 75% of structures according to the IMGT numbering scheme are shown. In each heatmap, the start and end positions of CDR1, 2 and 3 are highlighted. **D)** Inter-chain contact maps for 4 TCRs known to bind MHC with reversed orientation. The distance (Å, colour bar) is calculated between all pairs of atoms in the two residues, and the minimum distance is shown.

**Figure S11:**
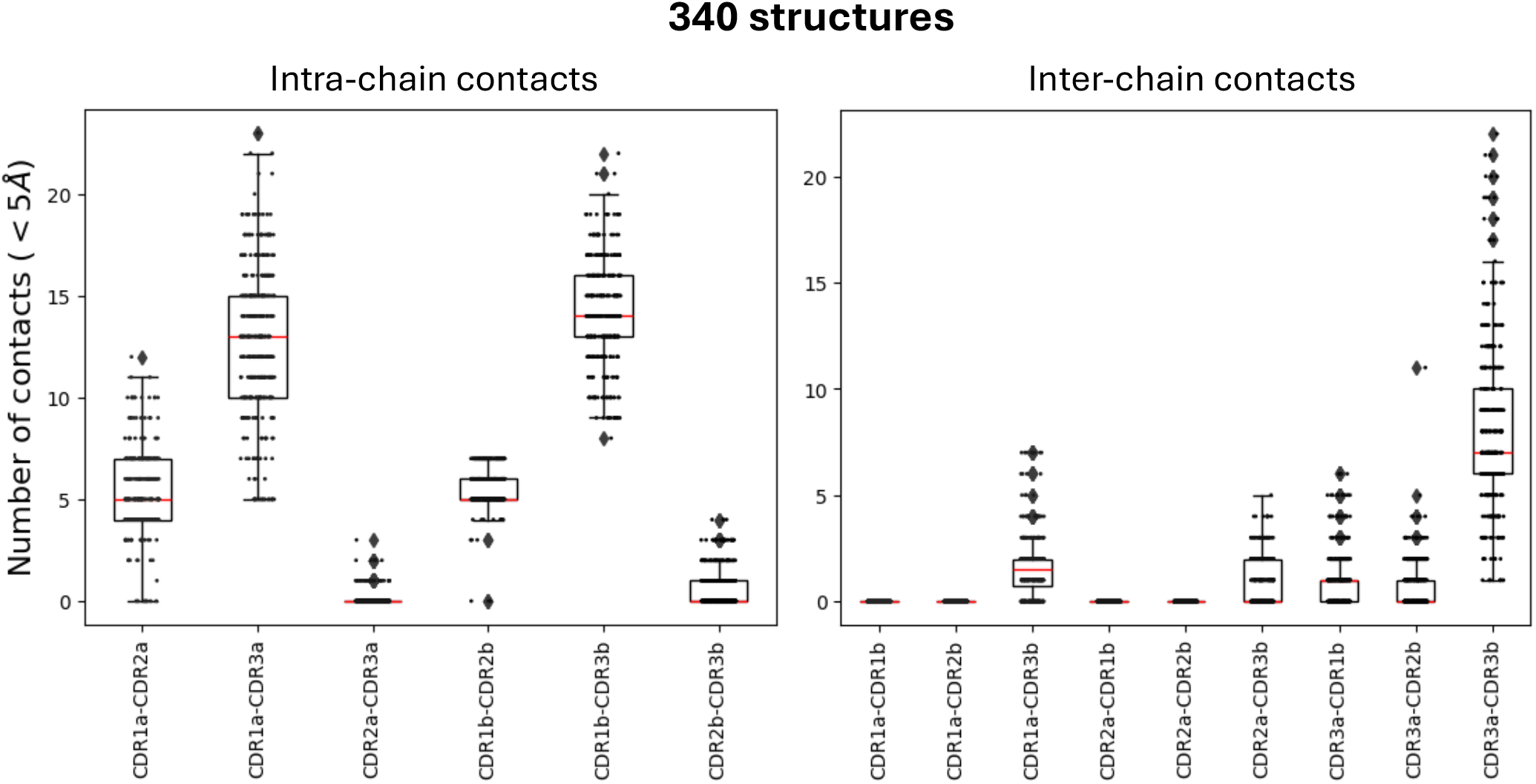
Number of contacts observed between CDRs. The number of intra- (left) and inter- (right) chain contacts observed in each of 340 TCR structures analysed is shown. A contact is defined as minimum distance between residues *<* 5Å. Red lines within the boxplots represent the median, and the box limits represent the quartiles. Whiskers of the boxplots extend to 1.5 the interquartile range of the lower or upper quartile.

**Figure S12:**
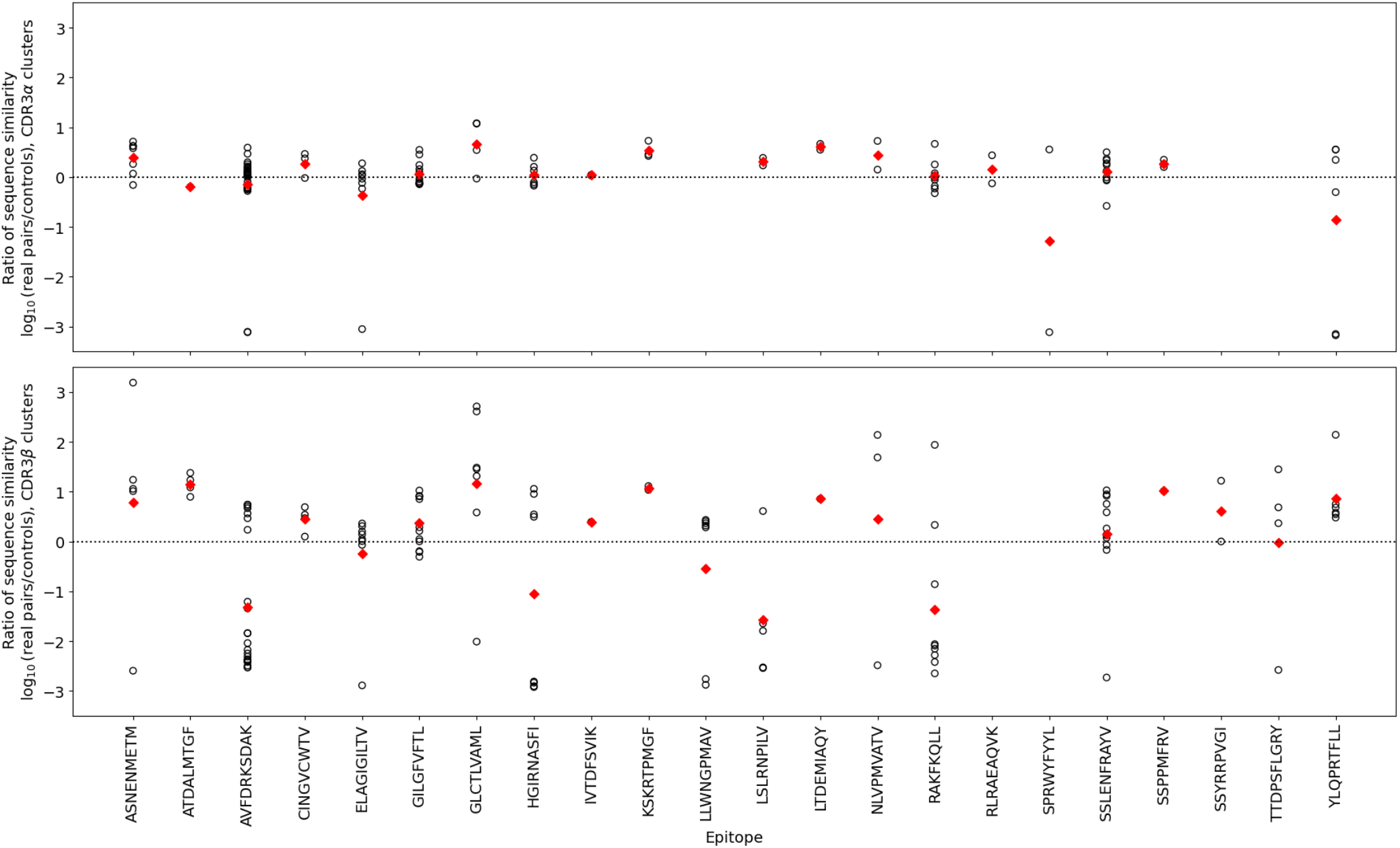
Ratio of sequence similarity between sequences paired to real clusters and random samples from epitope repertoires. For each cluster (of CDR3*α*, top and of CDR3*β*, bottom) in each epitope repertoire, the similarity between the paired CDR3s to the sequences in each cluster is calculated. 100 random samples of the same size from the same epitope-specific repertoire are also taken and their similarity calculated. The ratio between the median sequence similarity of the real pairs and of the random controls is shown (black circles, average median control similarity taken over all controls, ratio shown as log_10_, 0s imputed as 10^*−*4^ to be able to calculate the ratio and the log_10_ of the ratio). The red diamonds show the average across all clusters for that epitope.

**Figure S13:**
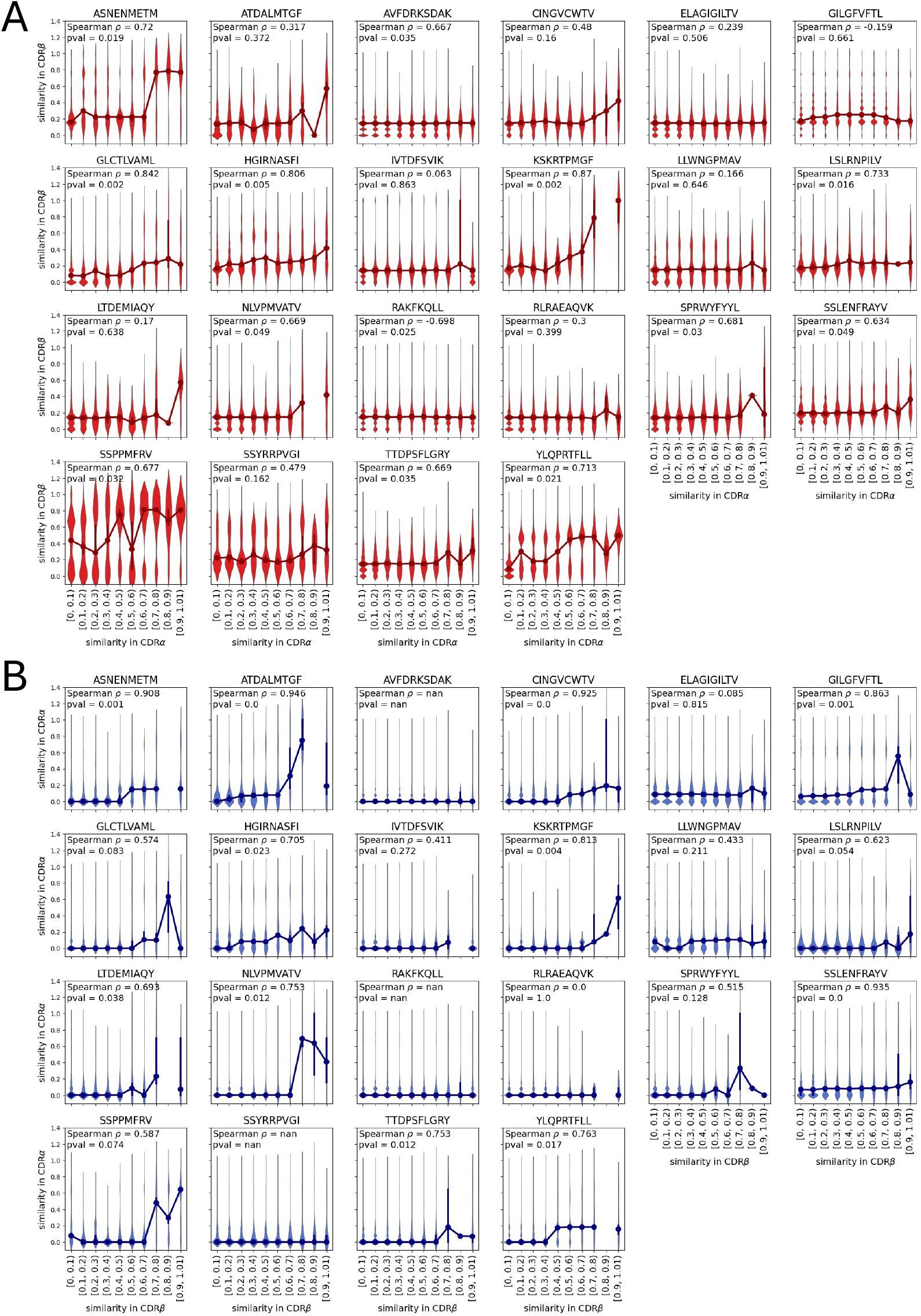
Correlation of CDR3 pairwise sequence similarity between the two chains. Pairwise similarity was calculated between all CDR3*α* and all CDR3*β* annotated for each epitope (including duplicate sequences). The pairwise similarity on the CDR3*α* was then binned and the similarity of the paired CDR3*β* examined (**A**, vice versa in **B**). The large circles joined by a solid line represent the median for each distribution, whilst the violin plots show the distribution of the underlying distances. Spearman correlation was calculated using the mid-point of each bin, and correlating with the median of the values for that bin.

**Figure S14:**
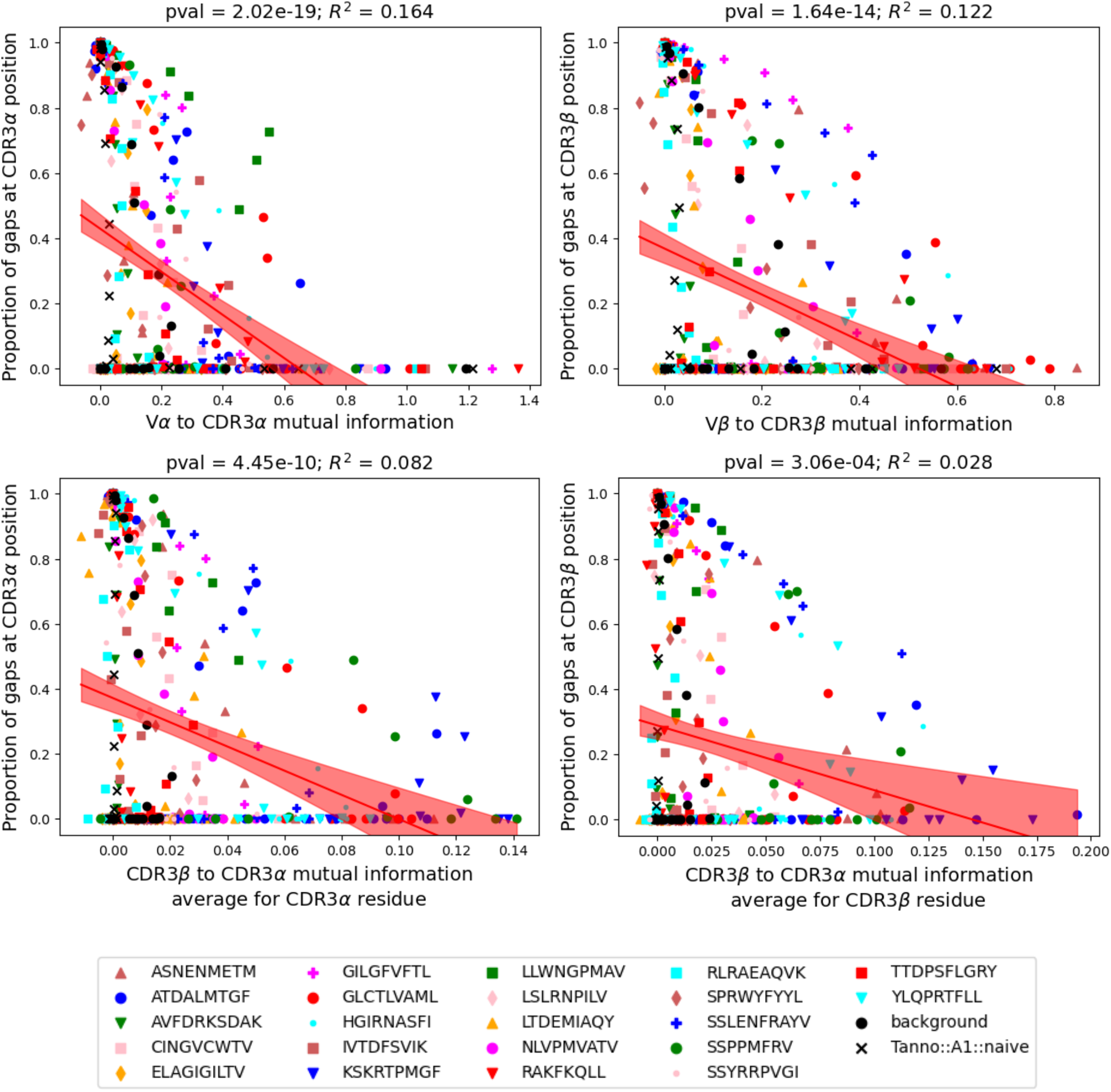
Contribution of gaps to MI information. Average MI per CDR3 (*α* on the left, *β* on the right) residue with V gene (top) or CDR3 (bottom). pval and *R*^2^ of shown fit are reported. The shaded area represents the 95% CI of the fit.

**Figure S15:**
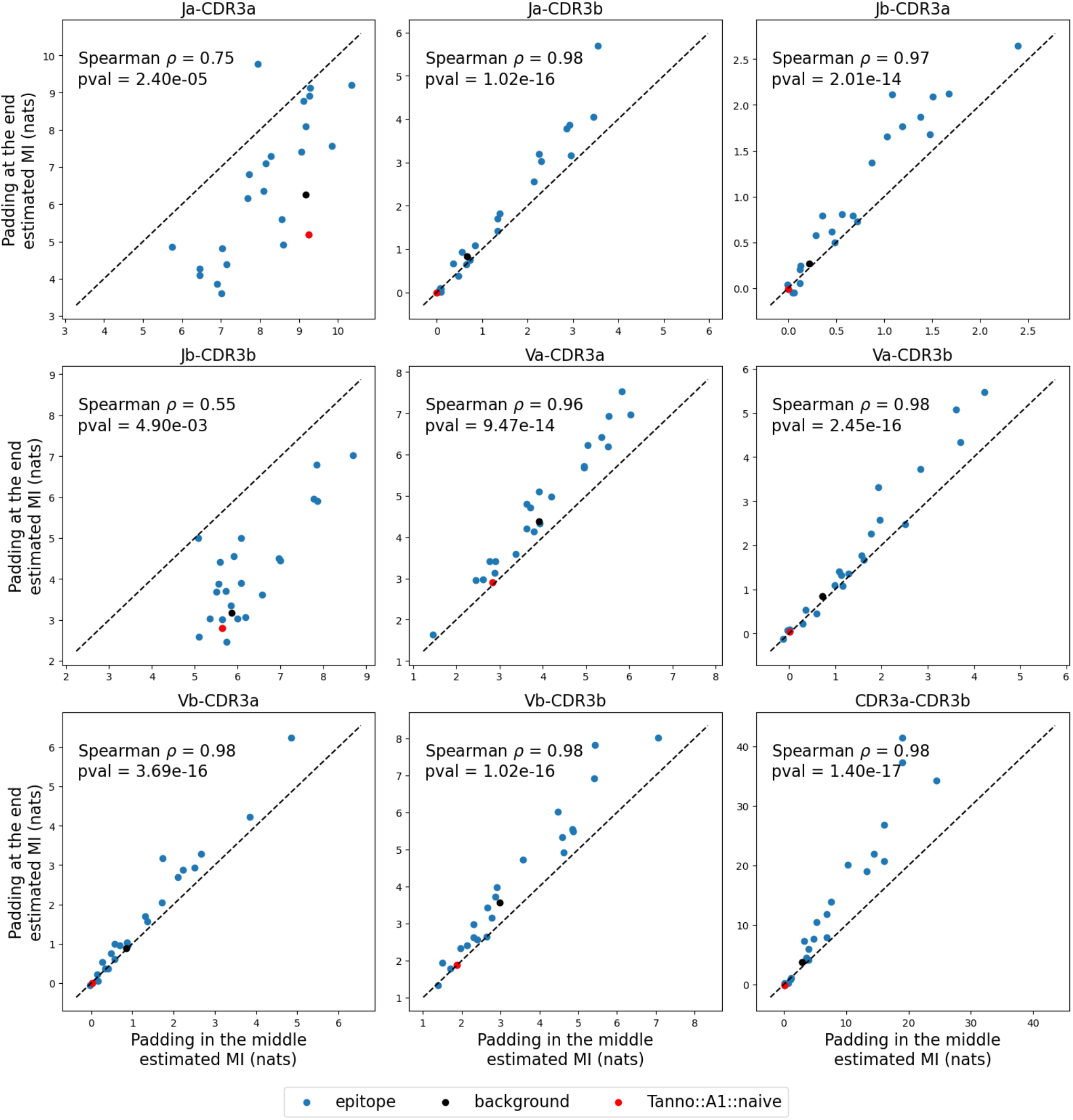
Comparison of estimated MI when different padding methods are used. For each pair of variables, the comparison of the MI estimated on CDR3s with padding in the middle (position 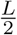 where *L* is the length of the CDR3 sequence) or at the end of each sequence is shown. The *y* = *x* diagonal is shown with a dashed line. Only pairs of variables including the CDR3 are shown in the plot, as V and J MI estimation is not affected by the padding strategy. Spearman *ρ* and p-value for each correlation are reported.

**Figure S16:**
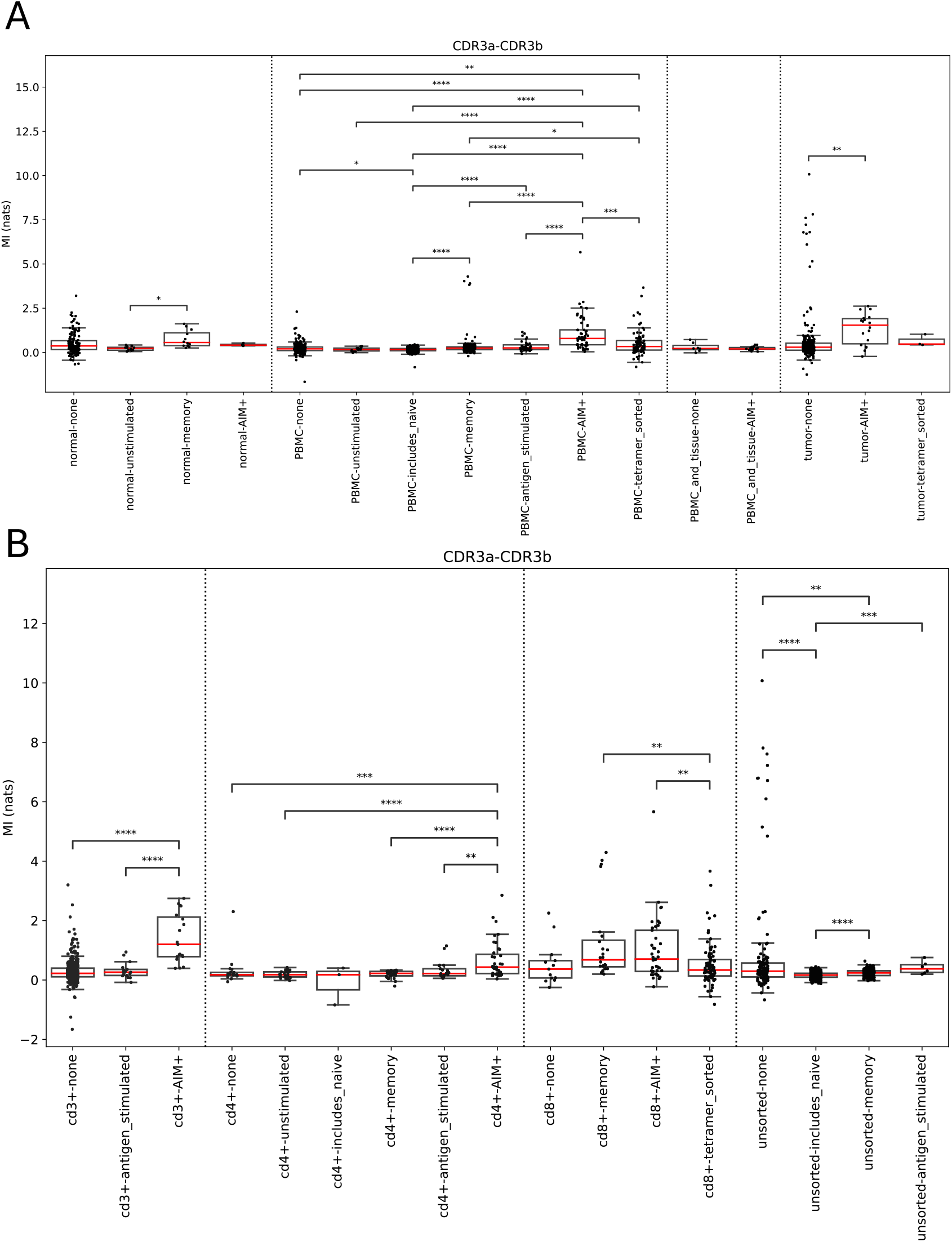
MI of datasets from OTS database. Calculated MI is shown for all included studies in the OTS databse, grouped according to the provided TSource and TSubtype (**A**) or the provided TType and TSubtype (**B**). The categories from the database are further grouped to give more meaningful larger groups for the comparisons. The edges of each boxplot show the quartiles, and the whiskers extend to 1.5 the interquartile range. Dotted lines separate comparison classes for the p-values. Mann-Whitney p-values with Benjamini-Hochberg correction are calculated within each class. Significant p-values are indicated by stars: **** < 10^*−*4^, *** < 10^*−*3^, ** < 10^*−*2^, * < 0.05

**Figure S17:**
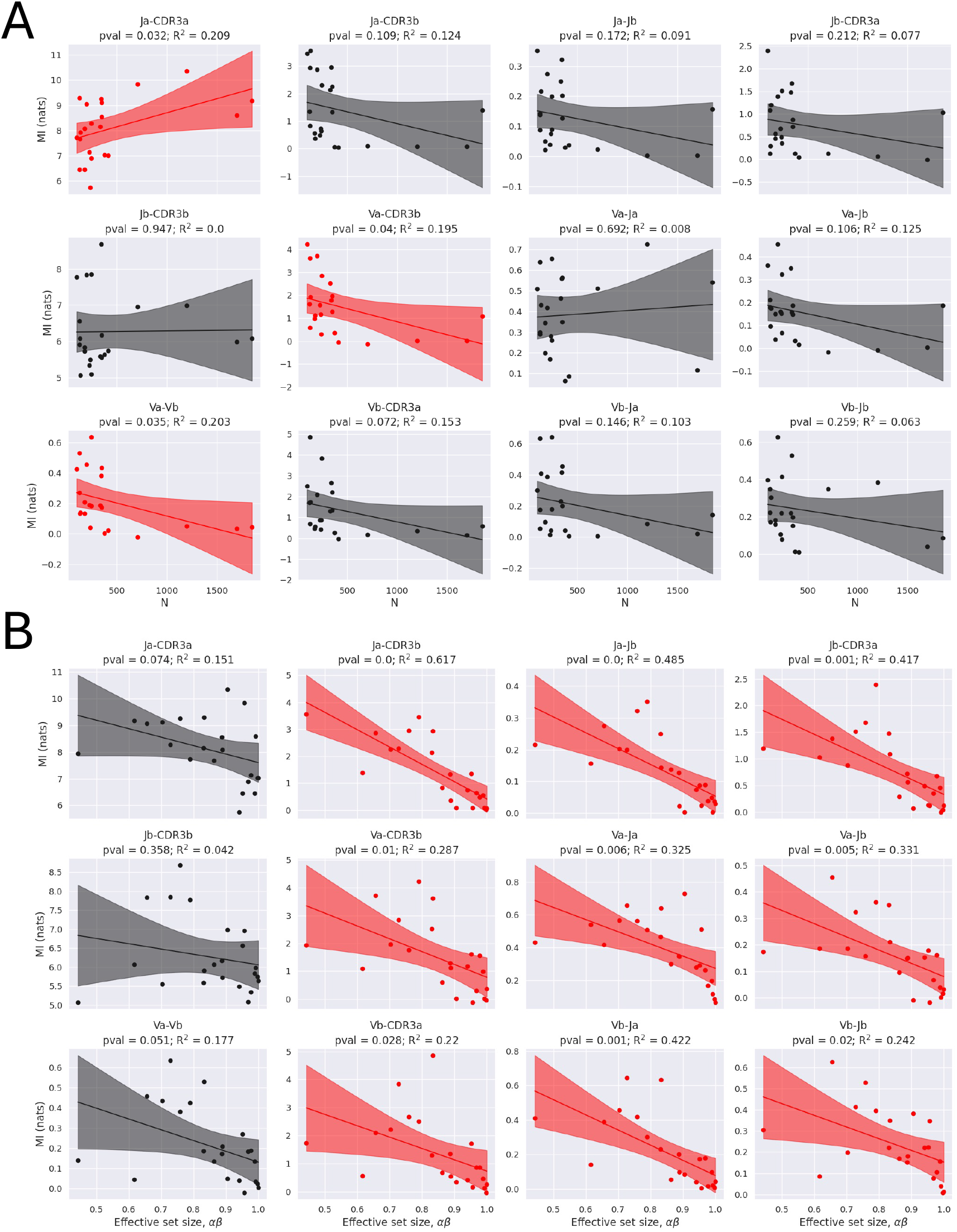
Correlation between mutual information and epitope repertoire size and mutual information and effective set size. For each epitope and each pair of variables the mutual information (MI) was estimated, correcting for finite size effects (see Methods). The correlation between MI and epitope repertoire size (N, **A**) or *αβ* effective set size (**B**) is shown for each pair of variables. Effective set size was calculated by grouping all sequences with Hamming distance *<* 3 on the paired chain, normalised by N. Correlation was calculated by fitting a linear regression. p-value for the F-test and *R*^2^ was calculated for each fit. Fits that have p-value<0.05 are highlighted in red. The shaded area represents the 95% CI of the fit.

**Figure S18:**
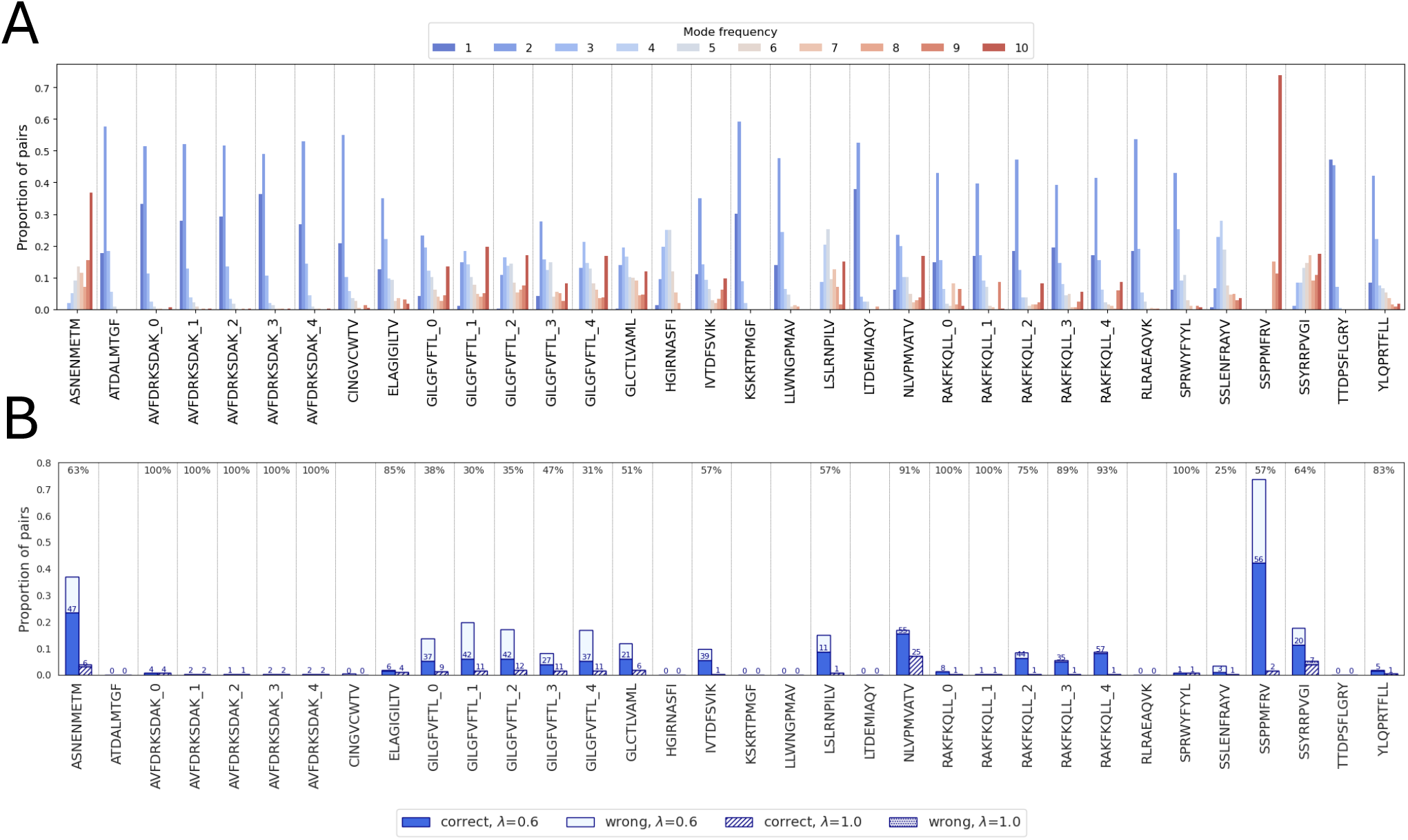
Stability of the pairs assigned by the MI-IPA. **A)** The bar chart shows the stability of the assignments made for each epitope. The x-axis shows the number of times an assignment is made across 10 repeats (i.e. the frequency of the mode, also colour-coded) and the y-axis shows the proportion of assignments that have that mode. The bar graph is repeated for each epitope, and for large epitopes all five subsamples are shown. The results shown correspond to pairing with *λ* = 0.6. **B)** The right-most bar for each epitope in **A** is broken down to show the proportion of stable assignments that are correct (with *λ* = 0.6 or *λ* = 1). The number on top of each bar indicates the number of correct pairs that were selected 10 times. The number at the top of each section indicates the percentage of correctly assigned pairs among pairs that are stably selected across repeats. Of note, some epitopes show stable pairs also in the chance expectation (*λ* = 1) scenario. These arise from individuals of size 1, i.e. examples where only a single *α* and a single *β* are available from an individual, which will always be assigned correctly independently of how well the model is learning.

**Figure S19:**
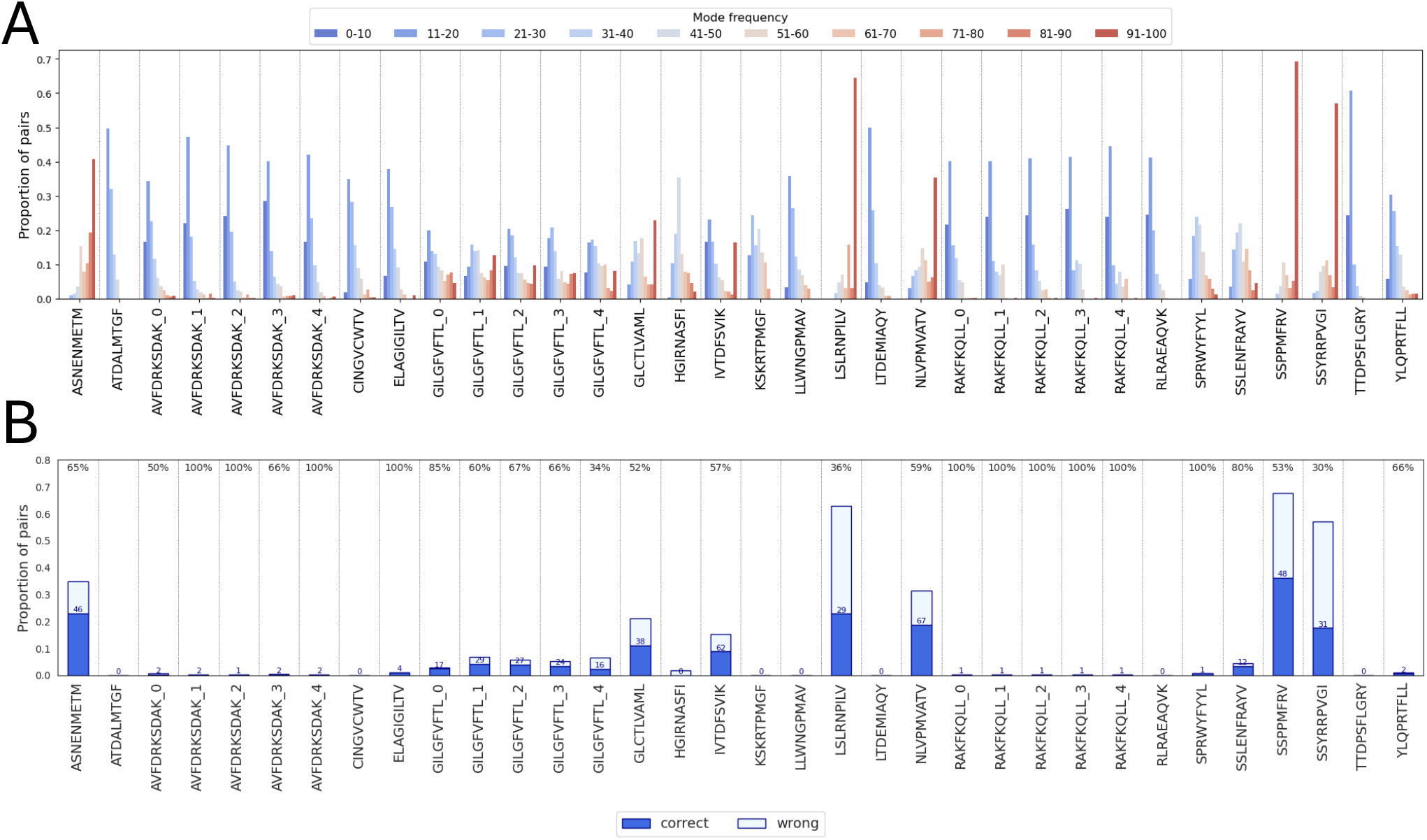
Stability of the pairs assigned by the GA. **A)** The bar chart shows the stability of the assignments made for each epitope. The x-axis shows the number of times an assignment is made across 100 repeats (i.e. the frequency of the mode, also colour-coded) and the y-axis shows the proportion of assignments that have that mode. The bar graph is repeated for each epitope, and for large epitopes all five subsamples are shown. The results shown correspond to GA run with *k* = 20 on Levenshtein distance. **B)** The right-most bar for each epitope in **A** is broken down to show the proportion of stable assignments that are correct. The number on top of each bar indicates the number of correct pairs that were selected *≥* 95 times. The number at the top of each section indicates the percentage of correctly assigned pairs among pairs that are stably selected across repeats.

**Figure S20:**
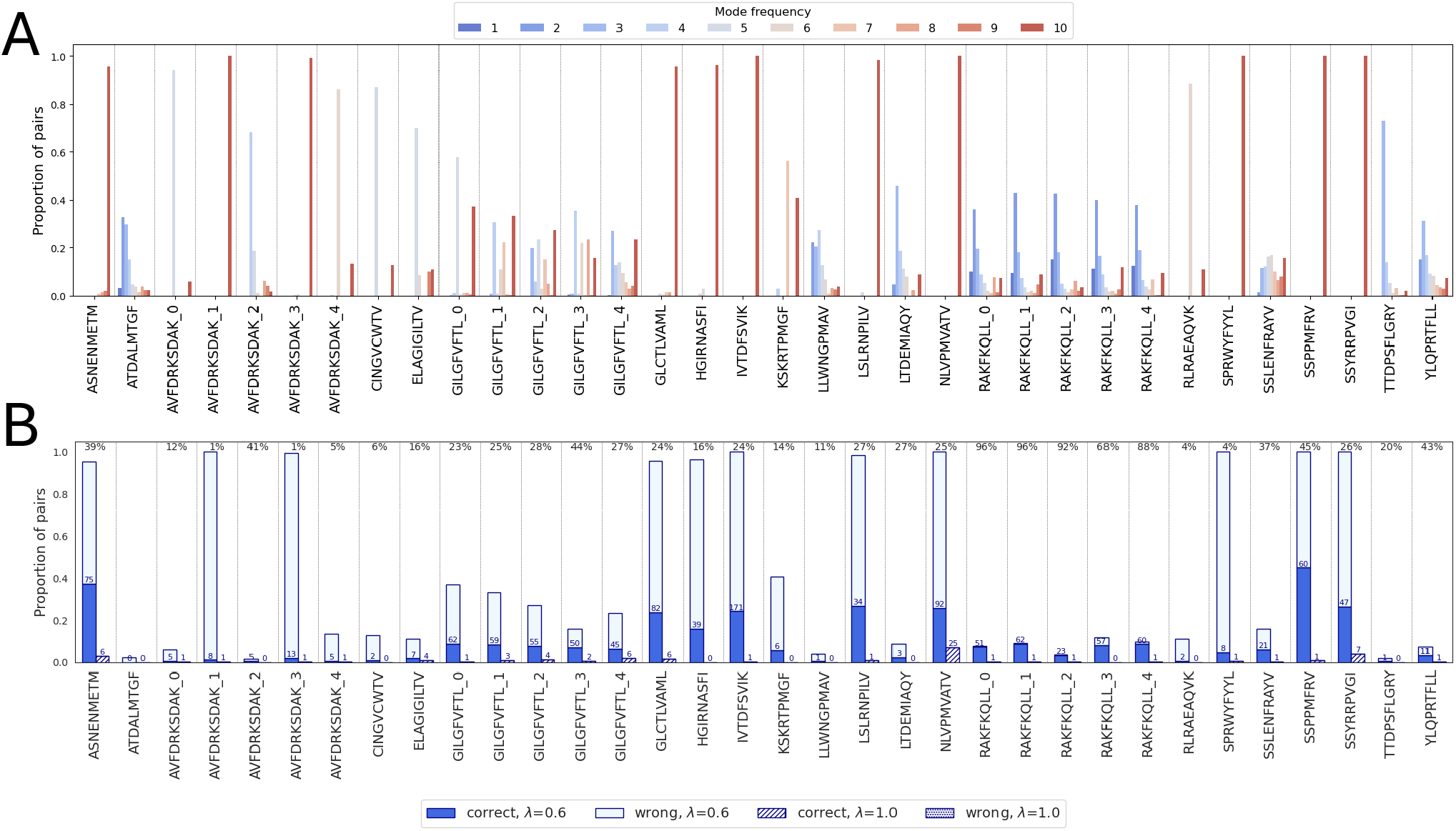
Stability of the pairs assigned by the GA+MI-IPA. **A)** The bar chart shows the stability of the assignments made for each epitope. The x-axis shows the number of times an assignment is made across 10 repeats (i.e. the frequency of the mode, also colour-coded) and the y-axis shows the proportion of assignments that have that mode. The bar graph is repeated for each epitope, and for large epitopes all five subsamples are shown (*λ* = 0.6, golden set from the GA, re-pairing allowed). **B)** The right-most bar for each epitope in **A** is broken down to show the proportion of stable assignments that are correct (MI-IPA run with *λ* = 0.6 or *λ* = 1, golden set from the GA, re-pairing allowed). The number on top of each bar indicates the number of correct pairs that were selected 10 times. The number at the top of each section indicates the percentage of correctly assigned pairs among pairs that are stably selected. Of note, some epitopes show stable pairs also in the chance expectation (*λ* = 1) scenario. These arise from individuals of size 1, i.e. examples where only a single *α* and a single *β* are available from an individual, which will always be assigned correctly independently of how well the model is learning.

**Figure S21:**
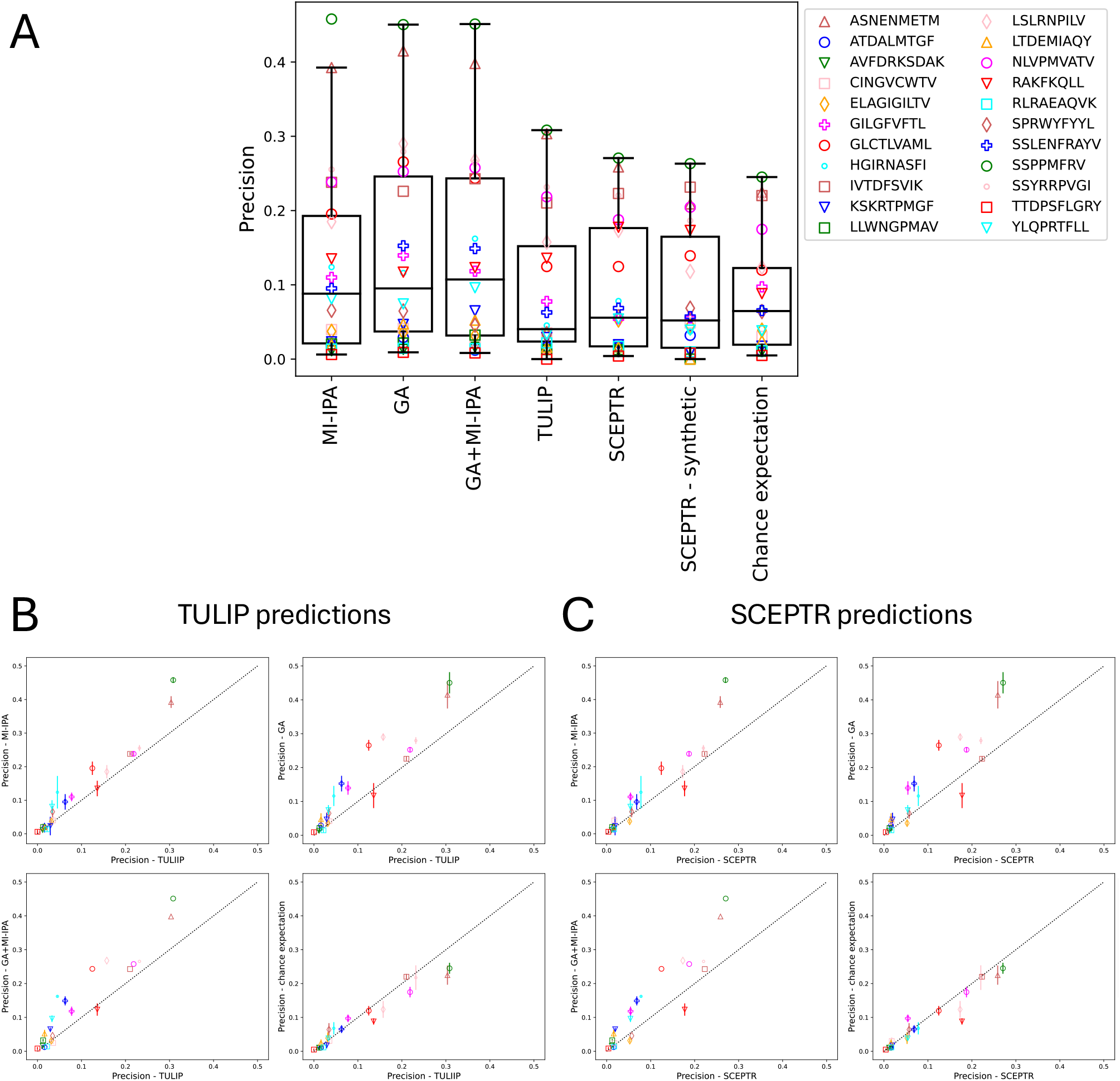
Benchmarking co-evolution-based pairing models against TULIP and SCEPTR. **A)** Precision for each epitope in the six models evaluated plus the chance expectation (MI-IPA with *λ* = 1). SCEPTR was run both with its default model (trained on paired data) and with a model trained on synthetic data, in which the pairing in the training set was random (SCEPTR - synthetic). The edges of each boxplot show the quartiles, and the whiskers extend to 1.5 the interquartile range. Precision of prediction based on the TULIP (**B**) or SCEPTR (**C**) output compared to assigned by MI-IPA, GA and GA+MI-IPA algorithms and chance expectation (MI-IPA with *λ* = 1). Error bars indicate the standard deviation over the repeats of the models for MI-IPA, GA and GA+MI-IPA. The dotted line represents the diagonal.

